# Antiviral immune response reveals host-specific virus infections in natural ant populations

**DOI:** 10.1101/2022.10.17.512467

**Authors:** Lumi Viljakainen, Matthias A. Fürst, Anna V. Grasse, Jaana Jurvansuu, Jinook Oh, Lassi Tolonen, Thomas Eder, Thomas Rattei, Sylvia Cremer

## Abstract

Hosts can carry many viruses in their bodies, but not all of them cause disease. We studied ants as a social host to determine both their overall viral repertoire and the subset of actively infecting viruses across natural populations of three subfamilies: the Argentine ant (*Linepithema humile*, Dolichoderinae), the invasive garden ant (*Lasius neglectus*, Formicinae) and the red ant (*Myrmica rubra*, Myrmicinae). We used a dual sequencing strategy to reconstruct complete virus genomes by RNA-seq and to simultaneously determine the small interfering RNAs (siRNAs) by small RNA sequencing (sRNA-seq), which constitute the host antiviral RNAi immune response. This approach led to the discovery of 41 novel viruses in ants and revealed a host-ant specific RNAi response (21 vs. 22 nt siRNAs) in the different ant species. The efficiency of the RNAi response (sRNA/RNA read count ratio) depended on the virus and the respective ant species, but not its population. Overall, we found the highest virus abundance and diversity per population in *Li. humile*, followed by *La. neglectus* and *M. rubra*. Argentine ants also shared a high proportion of viruses between populations, whilst overlap was nearly absent in *M. rubra*. Only a single of the total 59 viruses in our study caused active infection in more than one ant species, whilst six viruses infected one, but only contaminated another ant species. Disentangling active infection from contamination thus allowed us to show high host-specificity of active viral infections versus a decent degree of spillover of non-infecting viral contaminants across ant species, providing relevant information for ecosystem management.

## Introduction

Viruses and other pathogens are constantly exchanged between host individuals, be it by social interactions within species, predator-prey relationships between species, or by simply sharing a common environment (Boomsma et al., 2005; Fürst et al., 2014; French & Holmes 2020). Thus, the presence of pathogens inside or on a host’s body is not sufficient to determine whether it is a disease-causing agent of this host species, or whether it may only represent a contaminant in e.g. the host’s gut or on its skin. In addition to describing the variety of pathogens in a host, it is hence crucial to distinguish between active infections characterized by pathogen replication and a host immune response versus non-disease-causing contaminations, to determine the relevance of each pathogen for the disease dynamics and epidemiology within and across host populations.

The wide use of high-throughput RNA sequencing has recently started to provide unprecedented details on the variety of pathogens found in a host species. This includes so far less explored host-virus systems, exemplified by an extensive study of invertebrate viromes describing nearly 1500 new RNA viruses (Shi et al., 2016) by sequencing of viral RNA fragments (approx. 150 base pairs [bp] length) and assembly to the complete virus genomes. Whilst providing an excellent overview over the viruses found in a host, this does not, however, allow to distinguish whether these are real infections or only contaminants. To allow for this distinction, Webster and colleagues (Webster et al., 2015) have recently combined such long read sequencing with size-selected short read RNA sequencing to also recover the 21-22 nucleotides long small interfering RNAs (siRNAs) of viral sequence (Bernstein et al., 2001), which are produced as host antiviral RNA interference response (RNAi) response (Kemp & Imler, 2009).

In the RNAi response, the host immune system detects viruses by double-stranded RNA (dsRNA), which is either the infective viral stage (in double-stranded RNA viruses) or a replication intermediate (in single-stranded RNA viruses). The dsRNA is bound by the host enzyme Dicer and cleaved to small RNAs typically of sizes 21-22 nt. The diced dsRNA is then loaded to the Argonaute protein generating an RNA-induced silencing complex (RISC), where one of the strands is discarded while the other can then bind in a sequence-specific manner to more virus, leading to perpetuating cleavage to disarm viral replication (Wilson & Doudna, 2013). The host immune response against an infective and actively replicating virus therefore produces virus-sequence-specific siRNAs in the host’s cells. When Webster and colleagues used this combination of fragment size and sequence information in the well-studied *Drosophila melanogaster* system – currently the main model for studying individual immune responses in invertebrates (Buchon et al., 2014) – they were able to detect more than 20 novel viruses which represented active infections (Webster et al., 2015). Whilst highly abundant viruses are generally expected to represent host infection rather than contamination, this method is particularly powerful to define the infectivity of viruses found at low abundance.

In this study, we aimed to determine active viral infections that represent a disease threat and cause the RNAi host response in social insects, particularly ants. Like all social species, social insects face a high risk of transmission of infectious disease due to the close social contact between hosts (Cremer, et al., 2007; Schmid-Hempel 1998). Within the social insects (the social bees and wasps, the ants and the termites), viruses have been intensively studied in the honeybee and bumblebees, due to their ecological and economic importance as pollinators (for a review, see Grozinger & Flenniken, 2019). Bees are infected with a wide diversity of viruses. Transmission occurs either directly, mostly at flowers, which constitute shared food sources to many colonies and species – therefore often being ‘disease hotspots’– but also by worker drift to other colonies (Geffre et al., 2020), or via vectors, such as the *Varroa* mite (Boomsma et al., 2005; Chen et al., 2006). Bee viruses are hence transmitted within and among bee species (Alger et al. 2019; Fürst et al., 2014), as well as to other insects, including ants (Celle et al., 2008; Gruber et al., 2017; Levitt et al., 2013; Schläppi et al., 2020; Viljakainen et al., 2018). Viruses in bees may lie dormant or cause symptoms like deformed wings, paralysis or death, often causing important diseases such as the sacbrood disease in managed honeybee populations, but also the native bees and bumblebees, to which spillover regularly occurs in the field (Fürst et al., 2014; McMahon et al., 2015).

Ants and termites are less well studied, but we expect very different viral infection patterns than in bees due to their different ecology. They are highly territorial and feed exclusively within territories that they rigorously defend against neighbouring colonies, making cross-colony transmission very rare. Ants and termites are therefore expected to be less affected by viruses and bacteria, whose infectious stages can often only survive for a very limited period outside of a host in the environment (except if being able to produce long-lasting stages like spores, see Boomsma et al., 2005). Instead, infection of ants and termites frequently occurs via long-lasting infectious stages, picked up from the environment, such as bacterial and fungal spores from sporulating cadavers or from the soil (Boomsma et al., 2005; Cremer, Pull, & Fürst, 2018). Yet, as many ant species are scavengers and collect dead insects to feed their larvae, they can also pick up infections from virus-infected prey or when in vicinity to virus-infected other insects, like bee hives (Schläppi et al., 2020). Several studies on ants, most using molecular screening approaches, have reported that ants carry viruses (as found in 13 ant species, Supplemental Table 1; Baty et al., 2020), but determination which of them cause active infection is missing in most cases. Data on viral pathogenicity are so far only available for the well-studied invasive fire ant, *Solenopsis invicta*, where viruses may provide a useful tool in biocontrol (Manfredini et al., 2016; Valles & Hashimoto, 2009).

The social lifestyle of social insects has strong effects on disease dynamics. Colony members are typically highly related and live in dense communities where they frequently interact with each other to share information and food. These factors offer ideal conditions for pathogen transmission and persistence within a colony, but they also allow for coordinated and highly sophisticated cooperative disease defences, providing “social immunity” to the colony (Cremer et al., 2007; Cremer et al., 2018). Disease outbreaks are rare, due to efficient nest sanitation by antimicrobial compounds (Christe et al., 2003; Pull et al., 2018), active removal of disease vectors like the *Varroa* mite (Evans & Spivak, 2010), grooming of contaminated nestmates (Hughes et al., 2002; Rosengaus et al., 1998; Theis et al., 2015), removal and/or disinfection of infected brood (Pull et al., 2018; Rothenbuhler, 1964; Tragust et al., 2013) and changes in the social interaction network reducing its disease transmission properties after infection (Stroeymeyt et al., 2018). Social immunity is complemented by individual immunity of the insects, particularly their physiological immune system (Buchon et al., 2014; Konrad et al., 2012; Siva-Jothy et al., 2005), yet details on the expression and consistency of the RNAi response in ants is still missing, as is a comprehensive overview over their natural infection patterns.

To gain better insight into the viral infection patterns across ant species in the field, we studied one representative species each of the three major subfamilies of ants, the Argentine ant *Linepithema humile* (Dolichoderinae), the invasive garden ant *Lasius neglectus* (Formicinae) and the common red ant *Myrmica rubra* (Myrmicinae). For each ant species, we investigated RNA virus diversity, abundance and infectivity in three different populations to determine if infection patterns are species-specific or whether they follow regional patterns of infection. This choice of ants also allowed us to evaluate a possible effect of social structure on virus infections. All three studied ants are invasive species, yet we sampled only *Li. humile* (native to South America (Erickson, 1971) and *La. neglectus* (putative origin from the Black Sea area (Seifert, 2000)) from their introduced range in Southern Europe. Here, we also collected *M. rubra*. This species is native in these Eurasian populations from where it has been introduced to North America and Canada (Groden et al., 2005). Whilst forming small colonies with high territorial aggression in their native ranges, invasive ants form huge supercolonies in their introduced range (Giraud et al., 2002; Suarez et al., 2001; Ugelvig et al., 2008). Supercolonies are networks of nests, where the lack of aggression between individuals from the same supercolony enables constant exchange between nests and growth to enormous sizes (Cremer et al., 2008; Giraud et al., 2002). We expect these interactive networks to likely facilitate viral transmission across nest borders (Cremer, 2019; Ugelvig & Cremer, 2012). We applied the dual sequencing approach of long (∼ 150 bp) and short (∼ 30 bp) RNA sequencing on these ants, to determine their viral repertoire and overlap of active viral infections versus passive contaminants only, which has important implications for viral disease dynamics within and between natural populations of ants.

## Material and methods

### Ant collection

We collected the nonprotected ants, *Li. humile, La. neglectus* and *M. rubra* in April and May 2014. Each ant species was sampled from three different populations, two of which in relatively close geographic distance to one another in Spain and one, more distant to these, in Italy, resulting in a total of nine study populations (*Li. humile* from Orbetello, L’Escala and Sant Feliu de Guíxols; *La. neglectus* from Volterra, L’Escala and Seva; *M. rubra* from Monza, Ripoll and Vilallonga de Ter; Figure 1, details in Supplemental Table 2). After transport to the laboratory, we snap-froze ≥ 500 (mean 538, range 500-565) individuals per ant species and population in liquid nitrogen and stored the tubes at -80°C for further processing. Ant collection and all work in the laboratory followed European law and institutional guidelines.

**Figure 1.**
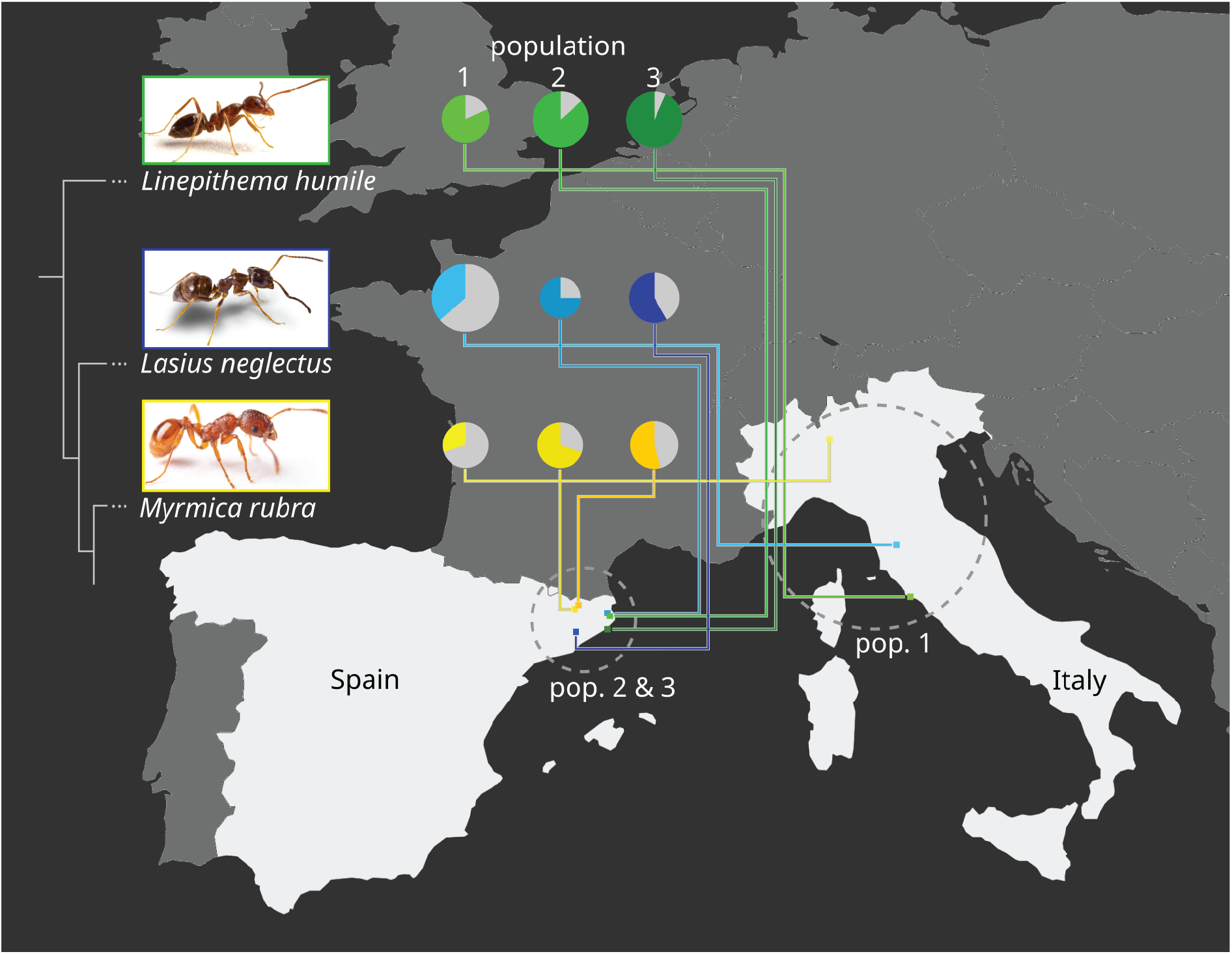
Ant host species and population viral diversity. We determined the viral repertoire of three populations each of three ant species, *Linepithema humile* (Dolicoderinae, green), *Lasius neglectus* (Formicinae, blue) and *Myrmica rubra* (Myrmicinae, yellow; phylogenetic relationship between the three subfamilies indicated by sketched tree). We sampled one population per species in Italy (pop. 1), and two in Spain (pop. 2&3; geographic relationship given in the map). Populations 1, 2 and 3 refer to Orbetello, L’Escala and Sant Feliu de Guíxols for *Li. humile*; Volterra, L’Escala and Seva for *La. neglectus*; and Monza, Ripoll and Vilallonga de Ter for *M. rubra*, respectively. For each population, the viral diversity (i.e. the number of different viruses detected) is indicated by pie size, and the proportion of actively infecting viruses in population-specific colour, as compared to only contaminating viruses (grey). Photocredit: Sina Metzler and Roland Ferrigato, ISTA.

### RNA extraction

Before starting the extractions, we removed the poison gland reservoir from all *La. neglectus* ants as the formic acid interferes with downstream applications. To obtain the highest RNA extraction efficiency possible, we processed ants in primary-pools optimized for ant number per species (5 ants each for *M. rubra* and *La. neglectus*, 10 ants for *Li. humile*). Primary-pools of ants were homogenized with 2 ceramic beads (2.8mm) in 350 µl QIAzol Lysis Reagent (Qiagen) for 2×2min at 30Hz in a TissueLyser II (Qiagen). After homogenization, an extra 400 µl QIAazol Lysis Reagent (Qiagen) was added along with 150 µl chloroform. The aqueous phase containing the total RNA was used for further RNA extraction following the standard protocol of the Qiagen miRNeasy 96 kit manual. We eluted in 45 µl H_2_O and ran the elution step twice with the same H_2_O to improve RNA concentration (although at the cost of RNA yield). Primary-pools were merged to generate a final pool of approximately 500-550 individuals (adults and brood) per site/species combination (resulting in 9 final pools; for details on sample compositions see Supplemental Table 2). RNA yield and quality of the final pools were measured on a Bioanalyzer (Agilent) and a Qubit Fluorometer (Invitrogen).

### Sequencing

Pools were sent for cDNA library preparation and sequencing to Eurofins Genomics GmbH (Ebersberg, Germany). For each pool two libraries were sequenced: one library of physically normalised RNA, random primed cDNA 150 bp paired-end on a HiSeq 2500 v3 in rapid run mode and one library of PAGE size selected (19-30 bp) on a HiSeq 2000 v3 in an Illumina platform.

### Bioinformatics of RNA-seq data

All paired-end RNA-seq reads (approx. 150 bp length) were quality-controlled: adaptors were removed, low quality regions were trimmed and reads less than 36 nucleotides (nt) of length were removed using Trimmomatic with default parameters (Bolger et al., 2014). The reads were *in silico* normalised to a maximum sequencing depth of 50 and contigs were assembled using Trinity v2.6.6 (Grabherr et al., 2011). Trinity seems to be sensitive to sequence variation, which is frequently encountered in virus sequence data, and therefore we extended the Trinity-contigs using CAP3 (Huang & Madan, 1999) and IVA (Iterative Virus Assembler) (Hunt et al., 2015). Contigs of at least 1000 nt in length were used for BLASTX similarity search (Altschul et al., 1990) with an e-value threshold of 10^−10^ against RefSeq viral protein databases downloaded from the National Center for Biotechnology Information (NCBI Resource Coordinators, 2017) on May 15^th^ 2018 to identify contigs of viral origin. Open reading frames (ORFs) were extracted for the potential virus contigs using NCBI’s ORF finder (NCBI Resource Coordinators, 2017) and a separate BLASTX similarity search was carried out with each of the ORFs. BWA-MEM (Li & Durbin, 2009) was used for mapping the clean RNA-seq reads against the annotated virus contigs to get RPKM counts (reads per kilobase per million mapped reads), which normalises read counts for virus genome length and sample sequencing depth. We focused our analyses on RNA viruses since DNA viruses are typically much larger and more difficult to assemble reliably since not the whole genome is transcribed.

### Bioinformatics of small RNA sequencing (sRNA-seq)

Small RNA sequencing provides a mix of short RNA fragments originating from three main processes in the host, (i) the RNA interference pathway producing siRNA against active virus infection, (ii) the PIWI pathway producing piRNA in response to transposons and (iii) the miRNA pathway which functions in regulating the expression of nuclear transcripts. As we were only interested in the virus-derived siRNA, we needed to disentangle the small RNAs resulting from the different pathways by bioinformatic processes.

The RNA interference pathway (RNAi) is triggered by double-stranded RNA (dsRNA), which is an intermediate of almost every active virus infection (Son et al., 2015) and hence the universal virus marker inducing the antiviral innate immune response and the production of small interfering RNAs (siRNAs). The biogenesis of siRNA is carried out by Dicer-2 (Dcr-2) (Wilson & Doudna, 2013), which targets dsRNA and digests it into approximately 22nt long fragments (Bernstein et al., 2001). The siRNAs are further processed to serve as a template in the RISC complex, which specifically targets sequences complementary to the respective siRNA (guide strand incorporated in the RISC complex) and digests these sequences. This efficiently reduces the viral population in an insect host. During an active virus infection Dcr-2 is therefore generating a high amount of ∼22nt fragments which can be used as markers for an active viral infection (Webster et al., 2015). Not only can we determine an active infection, but the sequence information available from the siRNAs can also be used to identify the viral sequences responsible for the infection, by mapping 22nt siRNAs to the contigs derived from the long-read sequencing.

Another important defence pathway, the PIWI pathway, produces a population of small RNAs (piRNA). Although there is some overlap in the small RNA populations produced by both pathways, the majority of piRNAs (24nt to 35nt, Hirakata & Siomi, 2016) are slightly larger than the siRNAs (predominantly 21nt -22nt, Gammon & Mello, 2015a). As the PIWI pathway is mainly active in the germline to protect against transposable elements, it is of less relevance for our purpose. To focus our analysis on the RNAi rather than the PIWI pathway, we therefore excluded from further analysis contigs, which did not meet at least one of the two following criteria: (i) induction of host RNAi response based on normalized sRNA-seq read count (RPKM) per contig being higher than 10 while the proportion of 21-22nt siRNA being higher than 50% to disentangle it from piRNA, (ii) abundance, whereby the normalized RNA-seq read count (RPKM) per virus genome was higher than 10 to allow for detection of viruses that have evolved to evade the host RNAi response.

The single-end short-RNA sequencing (sRNA) reads were first trimmed of adapters and of reads less than 15 nt long using a perl script ‘sRNA_clean.pl’ included in the VirusDetect pipeline (Zheng et al., 2017). In addition, ribosomal RNAs (rRNA) were removed by aligning the reads against the SILVA rRNA database (Yilmaz et al., 2013) using Bowtie2 v2.3.4.2 (Langmead & Salzberg, 2012). The filtered reads were then used for virus identification using the VirusDetect pipeline (Zheng et al., 2017), which first maps the reads against known viruses. Here we used the insect virus reference database ‘vrl_Invertebrates_220_U97’ downloaded from ftp://bioinfo.bti.cornell.edu/pub/program/VirusDetect/virus_database/v220/U97/. In the VirusDetect pipeline, the mapping was followed by a reference-guided assembly of reads that matched the database virus sequences. The unmapped sRNA reads were assembled *de novo* to obtain contigs that were used for BLASTN and BLASTX searches against virus nucleotide and protein databases, respectively, to describe novel viruses. Finally, the filtered sRNA reads were mapped against undetermined contigs from BLAST searches to obtain the siRNA and piRNA size distribution for each of the contigs to help in the discovery of potentially novel viruses (Zheng et al., 2017).

The filtered sRNA-seq reads were also mapped against the newly assembled virus genomes using Bowtie (Langmead et al., 2009). A modified version of the R package ViRome (Watson et al., 2013) was used for visualizing the sRNA size distribution of the mapped reads (available at: https://github.com/Edert/viRome_ggplot2).

### Phylogenetic analysis

Phylogeny reconstruction was carried out based on RNA-dependent RNA polymerase (RdRP) amino acid sequences of each virus and its nine best BLASTP hits. If the virus sequence matched several database viruses common to another virus of this study, their phylogenies were estimated jointly. In addition, a separate phylogeny consisting of only ant-derived virus RdRP sequences was reconstructed. Alignments were generated using E-INS-I method in MAFFT v7.313 (Katoh & Standley, 2013) and trimmed using trimAl v1.2 (Capella-Gutiérrez et al., 2009). Amino acid substitution models were selected by using ProtTest 3 (Darriba et al., 2011) and unrooted phylogenies were reconstructed using PhyML v.3.0 with Nearest Neighbor Interchange (NNI) tree topology search operation and approximate likelihood branch supports based on approximate Bayes method (Guindon et al., 2010).

## Results

### Description of 41 novel RNA viruses

Our dual sequencing strategy of RNA-seq (150 bp reads) and sRNA-seq (19-30 bp reads) revealed a total of 59 viruses in our samples of the three study populations each from the ant species *La. neglectus, Li. humile* and *M. rubra* (Table 1, Figure 2). 31% (18/59) of these viruses were previously described based on Trinity-assembled contigs derived from the RNA-seq data (Table 1, Figure 2). The remaining 69% (41/59) shared less than 90% RdRP amino acid identity with their best hit in BLASTP search, and are hence described as newly discovered viruses in this study (Edgar et al., 2022). For approximately half of these novel RNA viruses (21/41) we could reconstruct complete genomes, whilst the remaining (20/41) could only be recovered partially, either having one or two incomplete ORFs or missing essential genes such as those encoding for a capsid or an RNA-dependent RNA-polymerase (RdRP). All these new virus identifications were based on the long sequence reads (RNA-seq). The sRNA-seq data, whilst essential for detecting the host antiviral response (see below), did not provide any additional information regarding novel virus sequences. The reason for this is that the assembly of the sRNA-seq reads to contigs yielded only fragmented information due to uneven distribution of the sRNA-seq reads on the virus genomes (for the mapping of the siRNA reads to the viral genomes that we obtained for both 14 known and 11 novel viruses see Supplemental Figure 1).

**Table 1.**
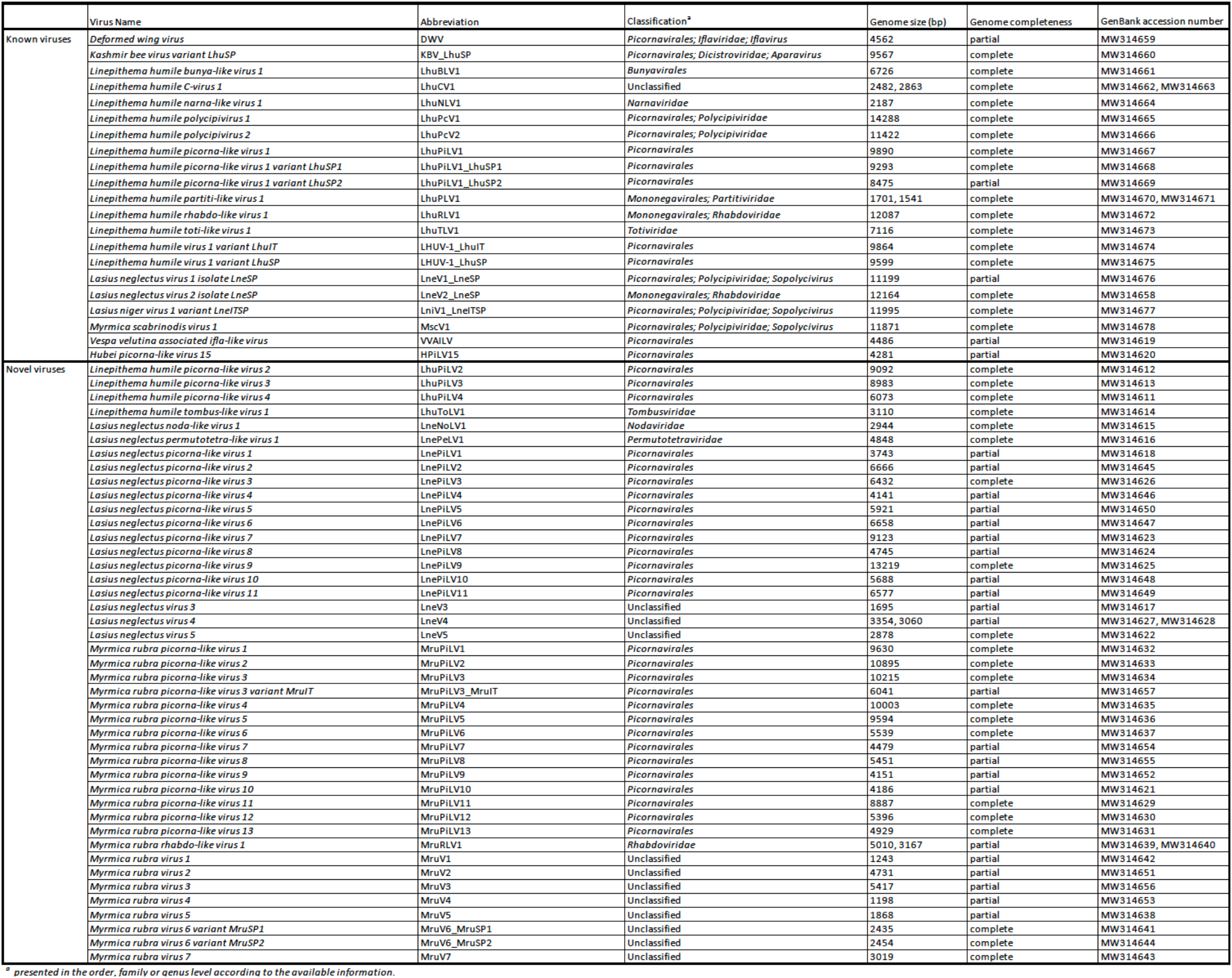
Previously described and novel viruses discovered in this study. Virus name, abbreviation used in this study and phylogenetic classification (to the degree of order, family or genus level depending on available information; unclassified in case the virus clustered with other unclassified sequences in the phylogeny or the virus genome assembly was missing RdRP) for all 59 viruses detected in this study. Note that some viruses were found as different variants (less than 95% similar at nucleotide level). Genome size and completeness based on RNA sequencing (RNA-seq), as well as GenBank accession number is provided (see Supplemental Table 1 for references on previously published ant viruses).

**Figure 2.**
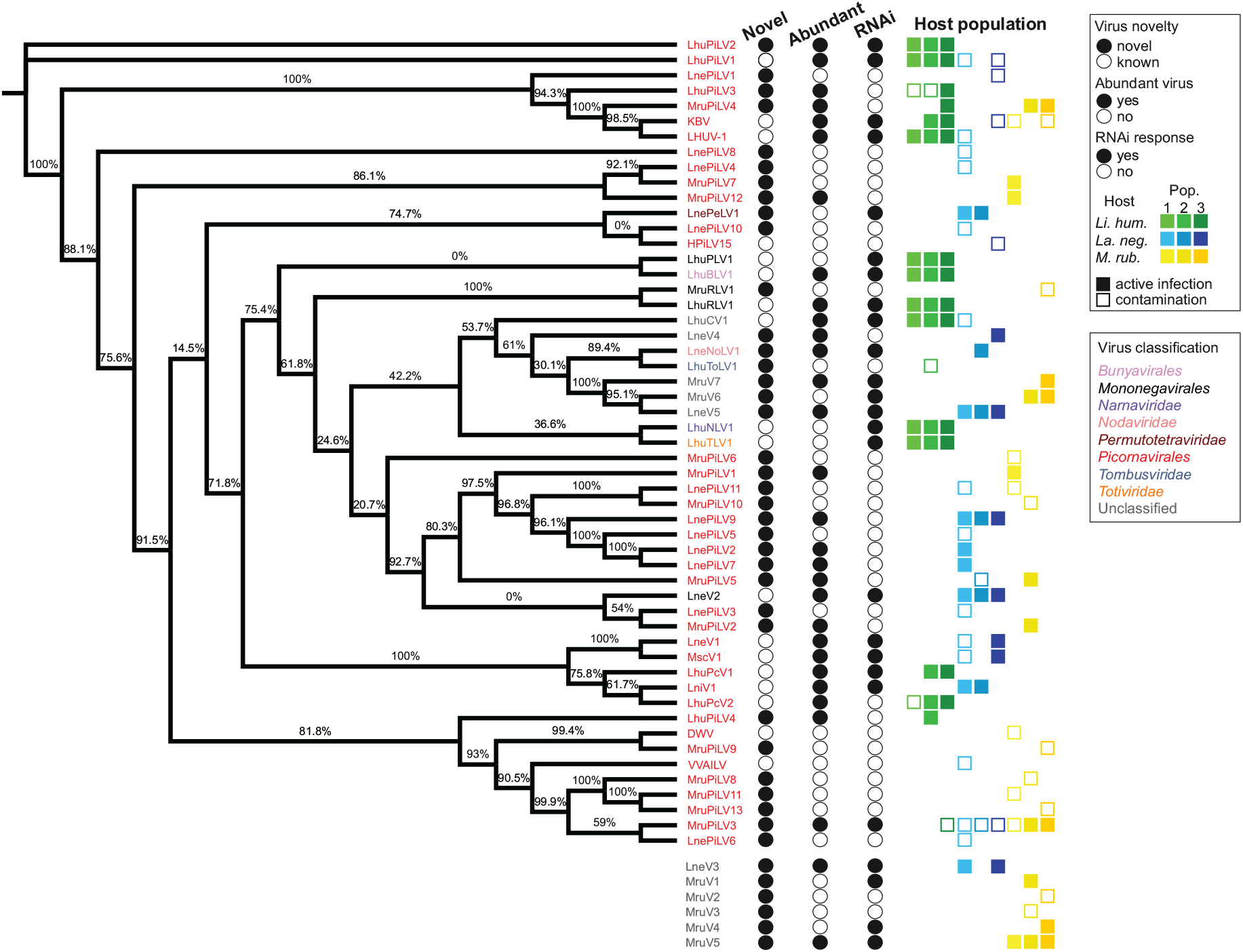
Virus phylogeny of the novel and known viruses. Phylogenetic position of the 53 viruses, for which we recovered the RdRP amino acid sequences (with the remaining 6 viruses, for which this sequence could not be determined shown below the tree) that we detected in the three populations each of our three studied ant species (*Li. humile* in shades of green, *La. neglectus* in shades of blue, *M. rubra* in shades of yellow). The topology of the tree is shown with posterior probabilities of tree branch supports based on approximate Bayes method in PhyML v. 3.0. Virus names are coloured according to their phylogenetic classification. Filled or empty circles indicate whether the virus is newly described in this study or already known, if it is abundant in at least one of the studied populations (defined by the normalized RNA-seq read count [RPKM] that maps to the virus genome being higher than 10), and whether any of the host populations has raised an RNAi response against the virus (defined by sRNA-seq RPKM higher than 10 while the proportion of 21-22nt siRNA is > 50%). All the host populations, in which the virus was detected are shown for each virus as a square, as a filled square in the population-specific colour (as in Figure 1), if consisting of an active infection, and an empty square where only representing a contamination (defined by the above abundance and siRNA criteria).

### Phylogeny of known and novel viruses

We had complete RdRP sequence information for 53 (out of the 59) viruses (16 known and 20 novel viruses with full genome information, as well as 15 out of 23 novel viruses with partial genome information). We first classified each of the 35 novel viruses with complete RdRP sequence separately by reconstructing a phylogeny together with its nine best BLASTP hits (Supplemental Figure 2). We then reconstructed a total phylogeny of all our novel and known 53 viruses with complete RdRP sequence (Figure 2). The majority of the branches had high support, but it is noteworthy that there were also a few branches with very low support and therefore the phylogeny remained partly unresolved. Nearly ¾ of all the viruses identified in our study (38/53) were classified as picorna-like viruses, which have non-enveloped positive-sense single-stranded RNA genomes typically encoding one large polyprotein (Zell et al., 2017). There were also novel viruses belonging one each to the *Permutotetraviridae, Nodaviridae, Rhabdoviridae* and *Tombusviridae* (Table 1). We were not able to classify 10 of the new virus sequences, as four of them clustered with unclassified viruses in the phylogeny (*La. neglectus:* LneV4 and LneV5, *M. rubra:* MruV6 and MruV7) and six could not even be included in the phylogeny due to their missing RdRP sequence (they were classified as viruses based on other identified virus proteins as follows: LneV3: coat/capsid protein; MruV1:capsid protein, MruV2: RNA helicase, MruV3: capsid, MruV4: capsid and MruV5: putative capsid).

### 2/3 of the viruses cause active infections in the ants

To separate contaminants (i.e. viruses that are not infective to the ants but are, for instance, ingested with the food) from viruses that actively infect the ants we set two requirements, either of which needed to be met: as actively infecting viruses multiply in the host, they are expected to either reach high abundances and/or to be fought by the host immune system, resulting in virus-specific siRNA. Note that viral abundance and host RNAi response can be independent classifiers of an active infection as some highly replicating viruses may suppress the immune response (Li, et al., 2002; Nayak et al., 2010; van Mierlo et al., 2012), or effective host siRNA prevents high viral loads from accumulating (Gammon & Mello, 2015b). We defined a virus as (i) abundant, when the normalized RNA-seq read count (RPKM) that maps to the virus genome was higher than 10; and (ii) capable of inducing the RNAi response, when the sRNA-seq RPKM was higher than 10 while the proportion of 21-22nt siRNA was higher than 50% to disentangle it from piRNA (see methods; Supplemental Table 3). Using these criteria, we found that only 63% of the viruses in our study (37/59) cause active infections, whilst 37% are likely to only be contaminants (Supplemental Table 3). 22 of the 37 active viruses were novel and 15 were already known (Figure 2). Hence, from the 18 previously described viruses, only three (the deformed wing virus (DWV), a well-known bee virus (Martin & Brettell, 2019; Wilfert et al., 2016); the Hubei picorna-like virus 15 (HPiLV15), identified from a mix of arthropods (Shi et al., 2016); the Vespa velutina associated ifla-like virus (VVILV), identified from the invasive hornet *Vespa velutina* (Dalmon et al., 2019)) were neither abundant nor did they induce an RNAi response. Of the remaining 15, the majority were both abundant and induced the RNAi response, whilst only three caused a raised host immune response despite not being abundant. This shows that previous work applying classical long RNA-seq had already revealed many abundant and hence important viruses of ants. For the novel viruses, the picture was more diverse: 7/22 were both abundant and induced RNAi, 11/22 were only found to be abundant (e.g. LhuPiLV3, LnePiLV9 and MruPiLV4), and the remaining four caused an RNAi response but were not abundant (e.g. LnePeLV1 and MruV1). Hence, the latter could only be detected as an active infection based on their siRNA signature.

### The RNAi response is host- and virus-specific

We could detect a host RNAi response mostly against abundant viruses, as 72% (18/25) of the RNAi responses were found against abundant, and only 7/25 against non-abundant viruses. This confirms that high viral load is often predictive of an active infection, which then also triggers a host immune response. Our data reveal, however, the importance of direct measures of the host response, as nearly 30% of the viruses causing a siRNA response would otherwise have remained undetected. The size distribution of the small RNAs varied between species: *Li. humile* viruses showed distributions peaking at 21-22 nt with either 21 nt or 22 nt RNAs predominating depending on the virus (Supplemental Figure 3), whereas *La. neglectus* and *M. rubra* viruses showed distributions with a peak only at 22 nt (Figure 3). The size distributions of the small RNAs also revealed that sRNA reads map relatively equally to positive and negative strands of the viruses. Moreover, in all ant species, we found one to two viruses that deviated from that general pattern. They either showed wide shoulders around major peaks at 21-22nt in a low sRNA read count, even if their RNA-seq read count was high in abundance (LhuPcV2 in *Li. humile*; MruPiLV4 and MruPiLV5 in *M. rubra*), or they had both a very high RNA-seq and sRNA read count with the sRNA size distribution not indicating a clear overrepresentation of a certain length (LnePiLV9 in *La. neglectus*). These viruses also showed a bias towards the positive strand in the size distributions of sRNAs (Supplemental Figure 4).

**Figure 3.**
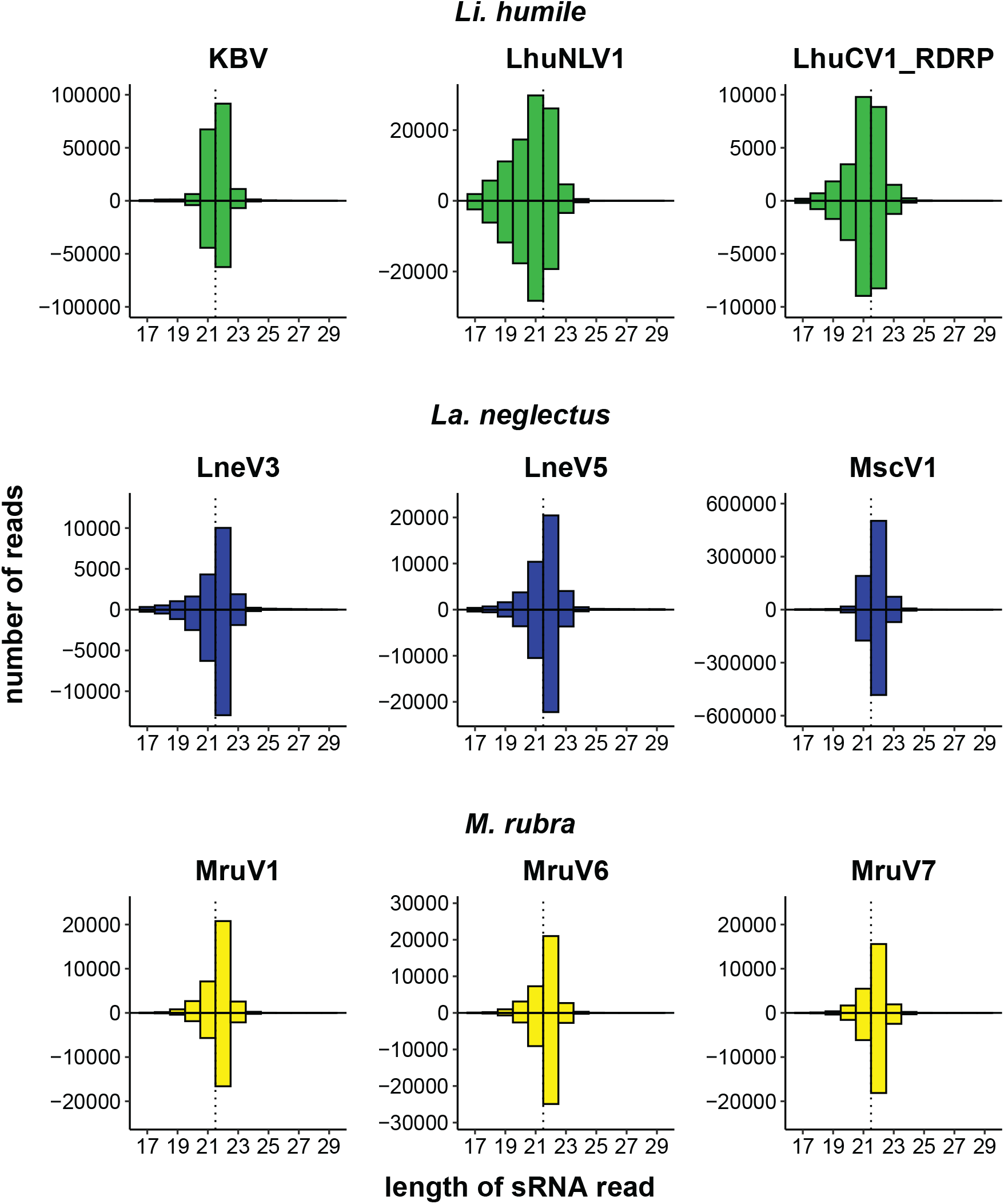
Host-dependent size distribution of the sRNA reads. For each ant species, *Li. humile* (green), *La. neglectus* (blue) and *M. rubra* (yellow), the three viruses with the highest number of sRNA reads are shown, separately for the positive strand (above the x-axis) and the negative strand (below the x-axis). Whilst *Li. humile* reacts with a virus-specific response of 21 nt or 22 nt (in 4 resp. 5 of the viruses, detailed in Supplemental Figure 3), both *La. neglectus* and *M. rubra* only produced 22 nt siRNA. Dotted vertical line separates 21 nt and 22 nt position. See supplemental Figure 4 for four viruses with atypical sRNA-seq read size distribution.

The efficiency of the host immune response, measured by the log-ratio of sRNA and RNA-seq RPKM values, showed considerable variation between viruses, indicating that some viruses generally elicit a higher RNAi response than others (such as LhuNLV1, LhuTLV1 and LhuPLV1 in *Li. humile* and MruV6 in *M. rubra*, Figure 4). Whilst the host response efficiency differed between viruses, it was highly consistent across host populations, which showed very little variation in their efficiencies towards the same viruses (Figure 4, Supplemental Table 3). The only exceptions to this were observed for LneV3 and LneV5 in the Volterra population of *La. neglectus*, where the efficiency was markedly higher than in the other populations of *La. neglectus* against the same virus.

**Figure 4.**
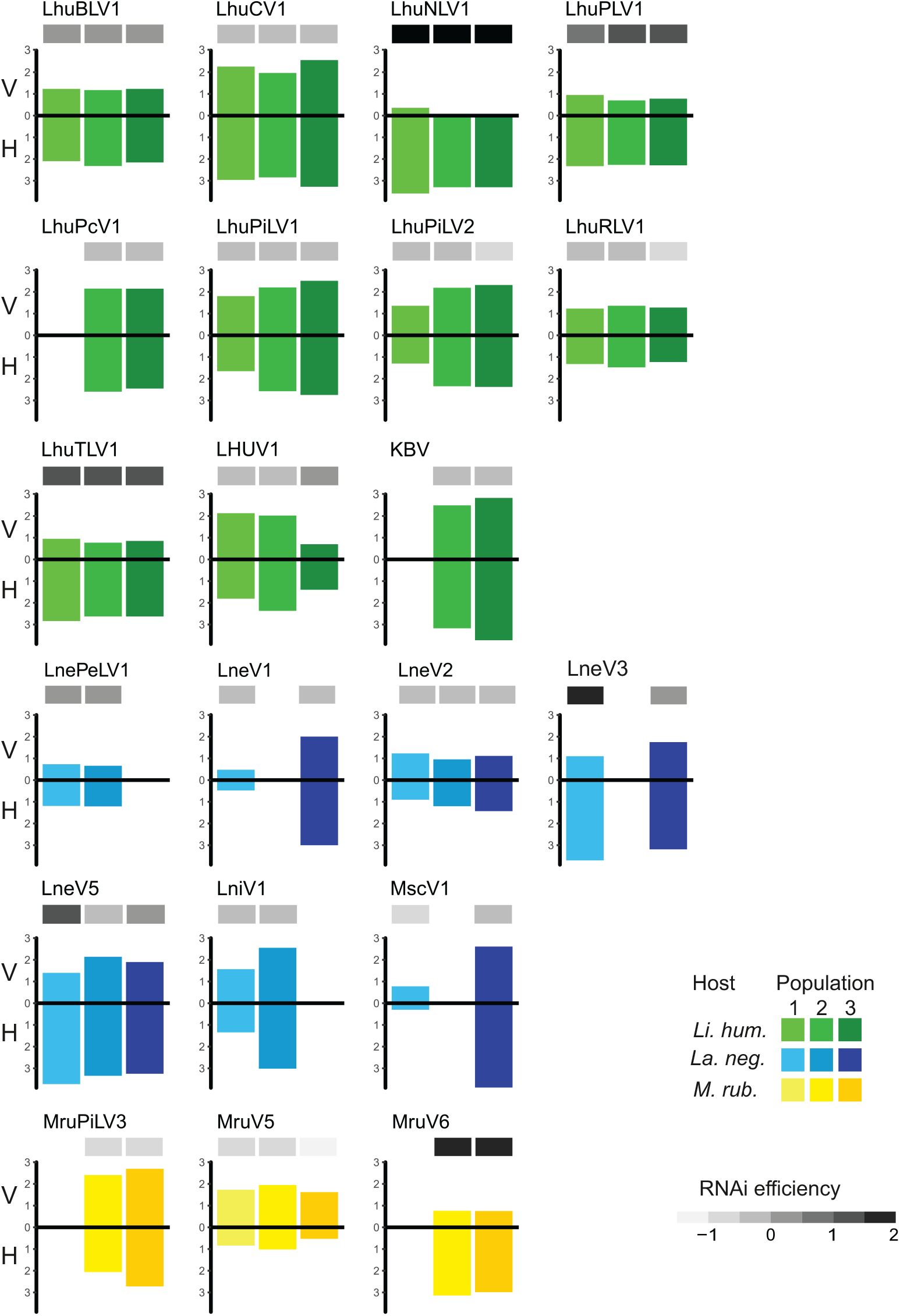
Efficiency of the host RNAi response. Virus load (V, measured as normalized RNA-seq read count [RPKM]; upper bar) and host RNAi response (H, measured as sRNA-seq RPKM; lower bar), as well as the resulting efficiency of the host RNAi response (ratio sRNA/viral load; the darker, the more efficient) is given for the 21 viruses that elicit an RNAi response in at least two study populations of the three ant species, *Li. humile* (in shades of green), *La. neglectus* (in shades of blue) and *M. rubra* (in shades of yellow); population-specific colour-coding as detailed in Figure 1.

### Viral infections differ in abundance and diversity between ant species

The overall virus abundance of the actively infecting viruses differed greatly between ant species: *Li. humile* had the highest number of virus-derived sequence reads based on RNA-seq (average normalized read count 1168 RPKM in the three populations), followed by *La. neglectus* (760 RPKM) and *M. rubra* (541 RPKM) (Table 2). The virus abundancies also differed between populations within a species: for example, the load of the Orbetello population (458 RPKM) of *Li. humile* was only ¼ of that of the Sant Feliu de Guíxols population (1939 RPKM). In *La. neglectus* the highest load was found in L’Escala (935 RPKM) and the lowest in Volterra (434 RPKM). The Vilallonga de Ter population of *M. rubra* had the highest virus load (901 RPKM), Ripoll had markedly less (642 RPKM) and the Monza population the lowest (79 RPKM) (Table 2 and Supplemental Table 3). These differences were also reflected in the viral species diversity, both within and between populations of the ant species.

**Table 2.**
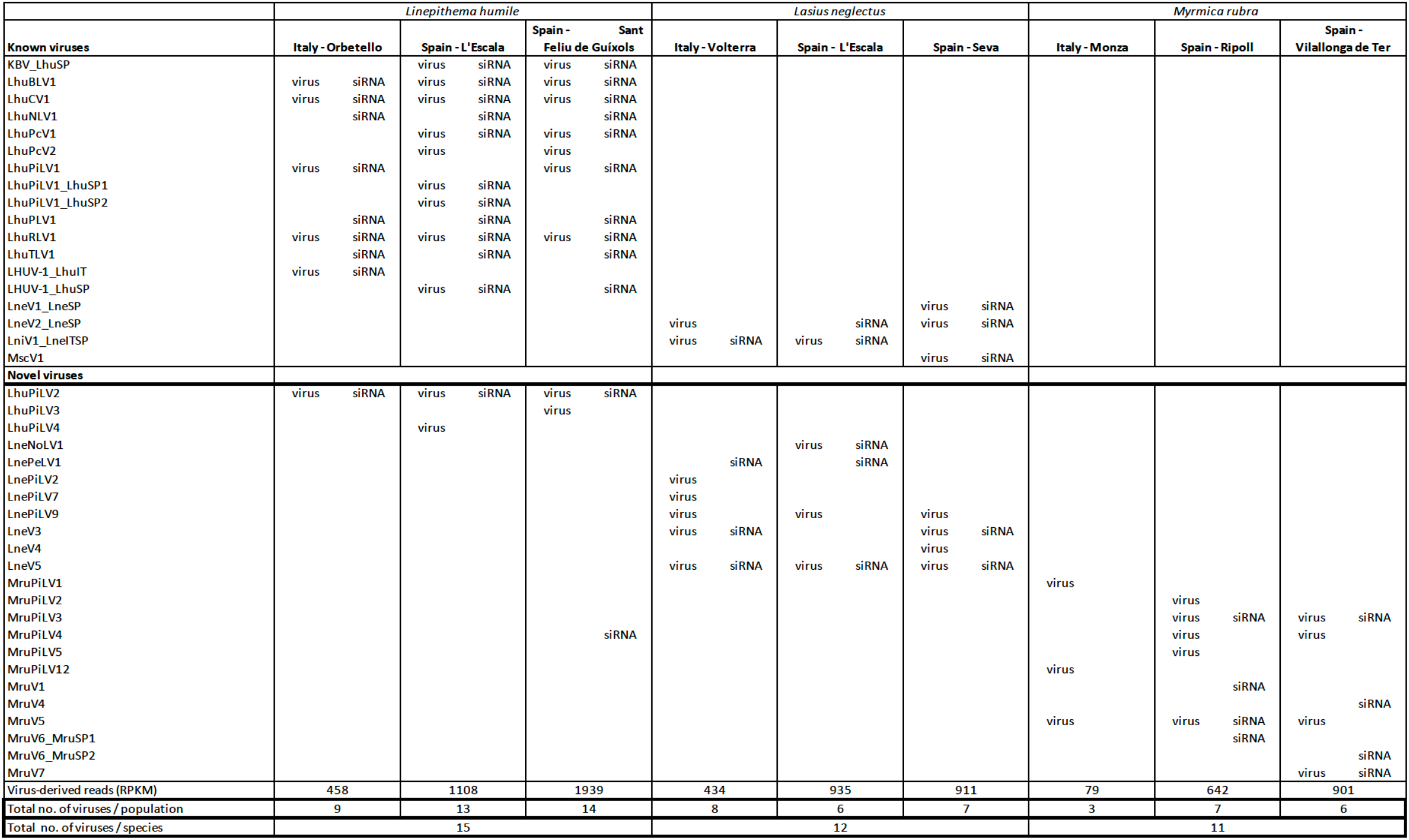
Actively infecting viruses in the three studied ant species. Abundant presence of viral sequences and presence of detectable levels of siRNA for the 37 actively infecting viruses for each ant species and study population, split up for already known and novel viruses. We identified a virus as causing an active infection in the host, when one of the two criteria were fulfilled: (i) viral abundance > 10 RPKM (based on RNA-seq, indicated by “virus”), (ii) sRNA RPKM > 10 and 50% of reads of sizes 21-22 nt (indicated by “siRNA”). For some viruses, both criteria were fulfilled at the same time, whilst others had either high RNA-seq or siRNA data (as detailed in the table). The total viral abundance per population (the sum of virus-derived reads; RPKM), as well as total number of viruses per population resp. species are given.

The highest diversity of actively infecting viruses was discovered in *Li. humile*, which contained a total of 15 distinct active viruses with a high within-population variation of 9 to 14 viruses. In *La. neglectus*, we found intermediate diversity with a total of 12 viruses (6 to 8 per population), whilst *M. rubra* had the lowest within-population diversity with only 3 to 7 out of the total 11 viruses found in each of the three populations (Figure 2, Table 2). We found the same pattern for between-population diversity: in *Li. humile*, 80% (12/15) of the viruses were shared between at least two, often even all three populations, whereas the proportion of shared viruses was only 50% (6/12) in *La. neglectus* and even only 36% (4/11) in *M. rubra* (Figure 5; Table 2).

**Figure 5.**
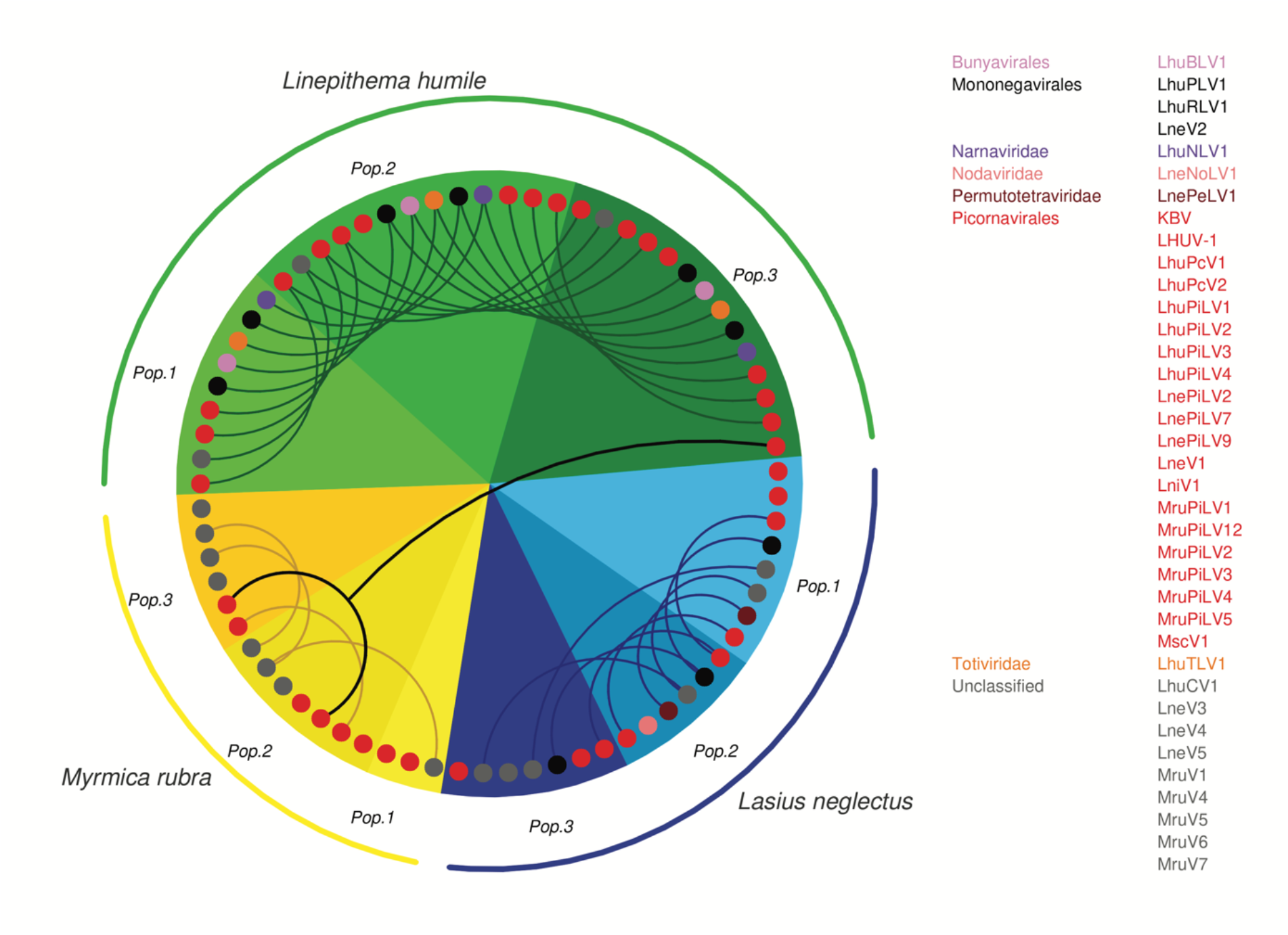
Presence and overlap of active viruses between ant species and populations. For each population of the three ant species, *Li. humile* (in shades of green), *La. neglectus* (in shades of blue) and *M. rubra* (in shades of yellow); colour coding of ant populations and the phylogenetic position of the actively infecting viruses presented as in Figures 1,2). Viruses occurring also in other populations are connected with a line, revealing a high, intermediate and low number of shared viruses between the populations of *Li. humile, La. neglectus* and *M. rubra*, respectively. Only a single virus (MruPiLV4) was shared between species, as it was found in two populations of *M. rubra* and one of *Li. humile*. Note that an interactive version of this figure is available under https://active-infections.science.ista.ac.at/.

The number of different viral classes found in each species were in line with the viral abundance and diversity. The actively infecting viruses harboured by *Li. humile* represented eight different viral classes (*Bunyavirales, Dicistroviridae, Narnaviridae, Partitiviridae, Picornavirales, Polycipiviridae Rhabdoviridae, Totiviridae*) and one unclassified virus. *La. neglectus* contained five virus classes (*Nodaviridae, Permutotetraviridae, Picornavirales, Polycipiviridae, Rhabdoviridae*) and three unclassified viruses, whereas six of the viruses found in *M. rubra* belonged only to one class, the *Picornavirales*, and five were unclassified.

Overall, this revealed that *Li. humile* was most affected by active viral infections, both in terms of viral species diversity and their abundance, whilst *La. neglectus* was intermediate and *M. rubra* least infected.

### Viral infections are specific to host species rather than region

Sampling each of our three ant species across a geographic area that overlapped between the three species allowed us to test if the viral infection patterns reflected host species-specificity or regional patterns. We clearly found that infective viruses were not geographically clustered but depended on host species across the sampled geographic range. Nearly all active virus infections were shared only between populations of the same ant species, but not across ant species, even if these were geographically very close in some sampling sites in Spain (Figures 1, 5). This host-specific infection pattern occurred even if the different populations were quite distant from one another (max. linear distance >700 km between populations, such as between *La. neglectus* populations Volterra and Seva) (Figure 1). Across species, only one virus (MruPiLV4, Figure 2, Table 2) was found to actively infect two of our ant species, namely one population of *Li. humile* and two populations of *M. rubra*, with the populations of the two species being 85 km apart (Figures 1, 5; Table 2; Supplemental Tables 2, 3). In addition, six more viruses caused active infection in one host species, and were also detected in low amounts and not inducing an RNAi response in other species (empty squares in Figure 2), suggesting it occurred as a potential contaminant in these other ant species. In the two closest populations, *Li. humile* and *La. neglectus*, both from L’Escala, even only a single virus was shared, which was a contaminant only for both host ants (MruPiLV3; Figure 2). We therefore could not detect any regional clusters of either infective or contaminating viruses, but instead found that viral infection was very specific to our studied ant species.

## Discussion

Viral infections of social insects have been studied extensively in bees (reviewed in Grozinger & Flenniken, 2019). In ants, however, only the invasive fire ant *Solenopsis invicta* was studied in detail (Hashimoto & Valles, 2008; Valles et al., 2004; Valles et al., 2007; Valles & Hashimoto, 2009; Valles & Rivers, 2019). We here used a novel dual approach for ants combining long and short RNA sequencing. This led to the description of 41 novel ant viruses, based on the long RNA-seq data. We also determined, which of the totally detected 59 viruses caused an active RNAi host response, based on the short sRNA reads mapping specifically to the viral genomes. Studying three populations each of three ant species representing the three major subfamilies of ants – *Linepithema humile*, Dolichoderinae; *Lasius neglectus*, Formicinae and *Myrmica rubra*, Myrmicinae – revealed host species-specific infection patterns with little regional or population-level effects, as well as host- and virus-specific efficiency of the RNAi response. Importantly, focusing on the actively infecting viruses – occurring either in high abundance or eliciting a detectable RNAi response – yielded different conclusions than a pure long read study would have revealed. Notably, our study suggests very little overlap of active viral infections across ant species, respectively subfamilies. If we had instead drawn our conclusions solely on viral presence based on long RNA reads, as used in most previous work, this would have led to a seven-fold overestimate of cross-species sharing of virus infections.

Following the establishment of this dual approach by Webster et al. in *Drosophila* (Webster et al., 2015), we took advantage of insect hosts responding to viral infection by the RNAi response, leading to the production of 21nt to 22nt long siRNAs that can be uniquely mapped to the viral genomes and are indicative of an active host immune response. The peak of the size distribution of the virus-derived sRNAs is known to be different across insect species. For example, in bees the size distribution shows a peak at 22 nt (Chejanovsky et al., 2014), whereas in *Drosophila melanogaster* and Culicine mosquitoes the distribution peaks at 21 nt (Göertz et al., 2019; Webster et al., 2015). Here we found that, in ants, there is no consistent peak size, but that the most abundant length depended both on the host ant and on the virus. Whilst the RNAi response in *La. neglectus* and *M. rubra* always peaked at 22 nt for all viruses, in *Li. humile* the most abundant lengths were both 21 and 22 nt, with the peak being either at 21 nt or at 22 nt, depending on the virus. Producing two major lengths of siRNAs within a single host species has also been observed for plants. In *Arabidopsis thaliana*, Dicer-like 4 (DCL4) produces 21 nt siRNAs and if DCL4 is suppressed by a viral suppressor of RNAi (VSR) then another enzyme (DCL2) is activated to produce 22 nt siRNAs (Deleris et al., 2006). Insect genomes typically encode for two Dicers, Dicer-1 and Dicer-2. In *Drosophila* the former has been associated with the generation of microRNAs that regulate host transcription, whereas only Dicer-2 is required for the production of siRNAs (Lee et al., 2004). The *Li. humile* genome also encodes for two Dicers, but it remains to be solved whether both are involved in the production of siRNAs, and whether this could explain the two sizes observed.

Applying this dual approach, we found 59 viruses in the natural populations of our three studied ant species, of which 18 were previously known RNA viruses and 41 were newly discovered by this study. Yet, only 63% (37/59) of these viruses caused active infections in the ants, using our criteria of either being highly abundant in their RNA-seq reads or eliciting an RNAi response. In approximately 50% of the cases, both criteria were fulfilled at the same time, whilst 32% (12/37) of the active viruses only showed high abundance and 19% (7/37) showed siRNA in the absence of a high viral load. This reveals that traditional approaches taking only viral abundance into account detect many important viruses but bear the risk of missing out on the ones that cause very efficient host response.

All three ants were able to raise an RNAi response, yet its effectiveness varied substantially against different viruses. Each ant species raised an extremely efficient response (as indicated by a high sRNA-seq/RNA-seq read count ratio) against several viruses (Figure 4). Whilst clearly infectious, these viral sequences can hence become too depleted to show high enough abundance to be picked up by long read analysis only. *Li. humile* and *La. neglectus* further contained three viruses that had both high viral load and high RNAi response, hence showing high viral replication despite the presence of a clearly raised host immune response. Interestingly, two of these viruses were first described in other species: KBV in the honeybee (Bailey & Woods, 1977; de Miranda et al., 2004) and LniV1 in the black garden ant *Lasius niger* (Olendraite et al., 2017).

On the other hand, some highly abundant viruses showed no detectable siRNA response, and four abundant viruses (LhuPcV2, LnePiLV9, MruPiLV4 and MruPiLV5) displayed atypical small RNA size distributions showing wide shoulders around the 22nt peak (Supplemental Figure 4). Intriguingly, Drosophila C virus (DCV) and Drosophila Nora virus have a similar atypical small RNA size distribution (in their case with a peak at 21 nt in their *Drosophila* host) (Webster et al., 2015). Both DCV and Nora virus are known to encode a viral suppressor of RNAi (VSR) (Lopez et al., 2018; van Rij et al., 2006), raising the question whether the abnormal sRNA size distribution found for the four abundant viruses not eliciting a normal RNAi response in the host in our study might be a signature of a potent viral suppressor. Further support for these viruses encoding a VSR is their sRNA distributions displaying a skew towards the positive strand (Supplemental Figure 4), which is observed also in several other viruses encoding VSRs (Brackney et al., 2009; Han et al., 2011; Sabin et al., 2013). However, annotation of the ORFs of these ant viruses with similarity searches failed to find homologous proteins with such a function necessitating further research to evaluate whether any of the ORFs might indeed encode for a viral suppressor.

In our study, the number of different viruses per host species, including also the non-infective contaminant viruses, ranged from 17 to 29, which is in line with previous studies of viruses in arthropod hosts, e.g. 20 in the fruit fly *D. melanogaster* (Webster et al., 2015), 18 in the invasive fly *Drosophila suzukii* (Medd et al., 2018), 9 in the tick *Ixodes ricinus* (Pettersson et al., 2017), and 31 in the honeybee *Apis mellifera* (Remnant et al., 2017). Notably, the actively infecting viruses only reached much lower numbers of 3-14 viruses per ant population, and the number of viruses per population strongly differed between the three ant host species. *Li. humile* populations each contained a high number of viruses, whereas each *M. rubra* population was infected with much fewer viruses, with the Monza population of *M. rubra* containing only three infective viruses (Figures 2, 5). The viral clade diversity was also high in *Li. humile* and *La. neglectus*, whereas all identified viruses of *M. rubra* belonged to picorna-like viruses (Figures 2, 5). As all the ant species showed a clear RNAi response against most of the highly abundant viruses, these differences are unlikely to derive from biological differences in the ant species’ ability to defeat virus infections. Instead, a more plausible explanation could be the connectivity of colonies by the movement of individuals between nests within a population since connectivity is expected to promote virus transmission and thus increase virus abundance as well as diversity. This explanation is supported by the red fire ant *S. invicta*, where larger, interconnected multiple-queen colonies harboured higher viral diversity and load compared to smaller single-queen colonies (Brahma et al., 2021). Whereas *M. rubra* is native in Italy and Spain (Groden et al., 2005), the sampled *Li. humile* and *La. neglectus* populations are invasive (Giraud et al., 2002; Ugelvig et al., 2008), forming supercolonies consisting of large networks of aggression-free nests (Holway & Suarez, 1999), which could explain the more extensive virus sharing in populations of *Li. humile* and *La. neglectus* than in *M. rubra*.

Similar to their differences of within-population viral diversity, the three ants also showed different levels of sharing of their viruses between their populations. The three *Li. humile* populations had high, *La. neglectus* an intermediate and *M. rubra* only very little overlap of viruses between populations. This is in line with the non-invasive *M. rubra* populations being independently established by local queen swarming. The study populations of the two invasive species, on the other hand, more likely originated from non-independent introductions by human-mediated dispersal (Cremer et al., 2008; Giraud et al., 2002; Holway & Suarez, 1999). Whilst the origin of *La. neglectus* is still putative and its invasion history only partially resolved, we know that it is a relatively young invader that was only detected in Europe in the 1970s, and whose invasive populations have a high potential to spread to new places (Seifert, 2000; Ugelvig et al., 2008). The Argentine ant, on the other hand, has a much longer invasion history and has established massive invasions around the world including in Europe for 120 years (Suarez et al., 2001). Interestingly, for one of our study populations (Sant Feliu de Guíxols), we could perform an across-years comparison to viral samples collected three years earlier (Viljakainen et al., 2018). This revealed that all 11 previously described viruses were still present in the population, supporting the notion that these viruses establish long-term relationships with their hosts. Ten of these virus species have also been found in the Argentine ant in New Zealand (Gruber et al., 2017) and in California (Viljakainen et al., 2018) implicating that either the viruses originate from a historical infection predating the worldwide invasion of the Argentine ant or that these viruses are transmitted across continents.

Insect viruses are usually able to infect various host species (McMenamin et al., 2018). We hence tested how many of the viruses were shared between the three ant species. One of our *a priori* hypotheses was that, if viruses may be able to use several ant species as hosts, we could find some geographic patterns across species. We could not find support for a regional viral infection pattern, not even in Spain, where the three ant species co-occur in a small geographic area with distances between the populations ranging from being less than one to max. 84 km (Figure 1). This is also in line with new populations of the two invasive ants establishing via human-mediated jump dispersal rather than small-scale dispersal by flight like the native species (Cremer et al., 2008; Giraud et al., 2002; Holway & Suarez, 1999). Instead, we discovered that the great majority of viruses in this study caused infection in a host species-specific manner, i.e. their infectivity pattern was not shared between the three ant species studied nor described earlier in any other insect. We only found a single actively infecting virus to be shared between *Li. humile* and *M. rubra* (Figure 5), leading to a cross-species sharing of only 1.7% (1/59). It is noteworthy, that a conventional study approach based only on long RNA reads would have overestimated this value seven-fold, as 11.8% (7/59) of the viruses were shared between ant species. In all six additional cases, however, we found the virus to actively infect only one of the species, whilst not fulfilling the criteria in the other species. This suggests that viral infectivity may be strongly host specific, whilst contamination occurs at a much higher frequency in ants.

Since the three study species belonged to three different subfamilies of ants, our resolution is not fine-grained enough to state whether the detected host specificity lies at the level of species or subfamilies. However, our data also suggest some cases of cross-infectivity between ant species and even subfamilies, as we found *La. neglectus* to be actively infected with LniV1 and MscV1, both described earlier from different ant species, either of the same genus (*Lasius niger*) or of a different subfamily (*Myrmica scabrinodis*, Myrmicinae) by Olendraite et al. 2017. As this study is based on RNA sequencing, it cannot be said for sure, however, if these viruses also caused active infections in the ants from which they were described. Moreover, *Li. humile* showed an active infection with the Kashmir bee virus (KBV) (Bailey & Woods, 1977; de Miranda et al., 2004) in this field study, which can also establish long term infections in laboratory colonies (Viljakainen et al., 2018). KBV is a well-known honeybee pathogen, that is also able to infect bumblebees (Singh et al., 2010) and wasps (Anderson, 1991). KBV has also been found to infect Argentine ants in New Zealand, where the viral load was markedly higher when the ant nests occurred close to honeybee hives (Gruber et al., 2017). This may also be true for some of the viruses that we have not identified as actively infecting in our study, maybe simply due to low prevalence among the pools of the 500 ants. One virus that has been shown to be a genuine ant-infecting virus is the Deformed Wing Virus (DWV), which has been observed in field-collected *M. rubra* at low levels, suggesting a spillover of this virus to ants, which is possible when ants nest close to beehives (Gruber et al., 2017; Schläppi et al., 2019). These observations support that some viruses can actively cross insect species barriers and actively infect different families (Formicidae and Apidae) within the order Hymenoptera, allowing for transmission from ant species to ant species or infection spillover from other hosts.

Viral transmission is particularly relevant in the light of some of these species being invasive pest species that may spread their diseases to the native ant fauna, similar to the well-described viral spillover from managed honeybees to the native bees and bumblebees (Fürst et al., 2014; McMahon et al., 2015). On the other hand, the potential exists for viruses to be used as effective biocontrol measures for invasive species (Oi et al., 2015; Oi et al., 2019), if their host specificity is narrow. Even if our study more than doubled the number of known ant viruses, we expect that our knowledge today represents only a minute fraction of the true viral diversity associated with the more than 15,000 ant species. To determine the specificity of infection and the transmission dynamics of viruses across the social insects, we advocate for more studies using the dual RNA-seq/sRNA-seq strategy to differentiate between active infections and non-disease-causing contaminations.

## Acknowledgements

We thank D.J. Obbard for sharing the details of the dual RNA-seq/sRNA-seq approach, S. Metzler and R. Ferrigato for the photographs (Figure 1), M. Konrad, B. Casillas-Perez, C.D. Pull and X. Espadaler for help with ant collection, and the Social Immunity Team at IST Austria, in particular J. Robb, A. Franschitz, E. Naderlinger, E. Dawson and B. Casillas-Perez for support and comments on the manuscript. The study was funded by the Austrian Science Fund (FWF; M02076-B25 to MAF) and the Academy of Finland (343022 to LV).

## Data Accessibility and Benefit-Sharing Statement

Ant collection was in compliance with European and national law. Benefits from this research accrue from the sharing of (i) our virus raw sequence data, submitted to NCBI under BioProject ID PRJNA681549, (ii) the assembled virus genomes, submitted to Genbank with accession numbers MW314611-MW314678, of (iii) providing detailed results on ant infection patterns (https://active-infections.science.ista.ac.at/) as well as (iv) providing tools for visualisation of sRNA data (https://github.com/Edert/viRome_ggplot2).

## Author contributions

SC and MAF designed the study; MAF coordinated the ant collection; AVG and MAF prepared the samples for RNA extraction; AVG performed the RNA extraction; MAF and TR developed the sequencing details with Eurofins; LV performed the bioinformatic analysis with input from JJ, LT, TE, TR and MAF; LV, JO, TE and SC conceived and prepared the figures with input from MAF; LV and SC wrote the manuscript with input from TE, AVG, MAF and TR. All authors approved the manuscript.

## References

Alger, S. A., Burnham, P. A., Boncristiani, H. F., & Brody, A. K. (2019). RNA virus spillover from managed honeybees (Apis mellifera) to wild bumblebees (Bombus spp.). PloS One, 14(6), e0217822.

Altschul, S. F., Gish, W., Miller, W., Myers, E. W., & Lipman, D. J. (1990). Basic local alignment search tool. Journal of Molecular Biology 215(3), 403–410.

Anderson, D. (1991). Kashmir bee virus--a relatively harmless virus of honey bee colonies. American Bee Journal, 131(12), 767–770.

Bailey, L., & Woods, R. (1977). Two more small RNA viruses from honey bees and further observations on sacbrood and acute bee-paralysis viruses. Journal of General Virology, 37(1), 175–182.

Baty, J. W., Bulgarella, M., Dobelmann, J., Felden, A., & Lester, P. J. (2020). Viruses and their effects in ants (Hymenoptera: Formicidae). Myrmecological News, 30, 213–228.

Bernstein, E., Caudy, A. A., Hammond, S. M., & Hannon, G. J. (2001). Role for a bidentate ribonuclease in the initiation step of RNA interference. Nature, 409(6818), 363.

Bolger, A. M., Lohse, M., & Usadel, B. (2014). Trimmomatic: A flexible trimmer for Illumina sequence data. Bioinformatics (Oxford, England), 30(15), 2114–2120.

Boomsma, J. J., Schmid-Hempel, P., & Hughes, W. O. H. (2005). Life histories and parasite pressure across the major groups of social insects. In M D. E. Fellowes, G. J. Holloway & J. Rolff (Eds.), Insect Evolutionary Ecology. (pp. 139–175). Wallingford: CABI.

Brackney, D. E., Beane, J. E., & Ebel, G. D. (2009). RNAi targeting of West Nile virus in mosquito midguts promotes virus diversification. PLoS Pathogens, 5(7), e1000502.

Brahma, A., Leon, G. L., Hernandez, G. L. & Wurm, Y. (2021). Larger, more connected societies of ants have a higher prevalence of viruses. Molecular Ecology 31, 859–865.

Brettell, L. E., Mordecai, G. J., Pachori, P., & Martin, S. J. (2017). Novel RNA virus genome discovered in ghost ants (Tapinoma melanocephalum) from Hawaii. Genome Announcements, 5(30), e00669–17.

Buchon, N., Silverman, N., & Cherry, S. (2014). Immunity in Drosophila melanogaster— from microbial recognition to whole-organism physiology. Nature Reviews Immunology, 14(12), 796.

Capella-Gutiérrez, S., Silla-Martínez, J. M., & Gabaldón, T. (2009). trimAl: A tool for automated alignment trimming in large-scale phylogenetic analyses. Bioinformatics, 25(15), 1972–1973.

Celle, O., Blanchard, P., Olivier, V., Schurr, F., Cougoule, N., Faucon, J., et al. (2008). Detection of chronic bee paralysis virus (CBPV) genome and its replicative RNA form in various hosts and possible ways of spread. Virus Research, 133(2), 280–284.

Chejanovsky, N., Ophir, R., Schwager, M. S., Slabezki, Y., Grossman, S., & Cox-Foster, D. (2014). Characterization of viral siRNA populations in honey bee colony collapse disorder. Virology, 454, 176–183.

Chen, Y., Evans, J., & Feldlaufer, M. (2006). Horizontal and vertical transmission of viruses in the honey bee, Apis mellifera. Journal of Invertebrate Pathology, 92(3), 152–159.

Christe, P., Oppliger, A., Bancalà, F., Castella, G., & Chapuisat, M. (2003). Evidence for collective medication in ants. Ecology Letters, 6(168), 19–22.

Cremer, S. (2019). Pathogens and disease defense of invasive ants. Current Opinion in Insect Science, 33, 63–68.

Cremer, S., Pull, C. D., & Fürst, M. A. (2018). Social immunity: Emergence and evolution of colony-level disease protection. Annual Review of Entomology, 63, 105–123.

Cremer, S., Armitage, S. A., & Schmid-Hempel, P. (2007). Social immunity. Current Biology, 17(16), R693–702.

Cremer, S., Ugelvig, L. V., Drijfhout, F. P., Schlick-Steiner, B. C., Steiner, F. M., Seifert, B., et al. (2008). The evolution of invasiveness in garden ants. PloS One, 3(12), e3838.

Dalmon, A., Gayral, P., Decante, D., Klopp, C., Bigot, D., Thomasson, M., Herniou, E. A., Alaux, C. & Le Conte, Y. (2019). Viruses in the Invasive Hornet Vespa velutina. Viruses 11(11), E1041.

Darriba, D., Taboada, G. L., Doallo, R., & Posada, D. (2011). ProtTest 3: Fast selection of best-fit models of protein evolution. Bioinformatics, 27(8), 1164–1165.

de Miranda, J. R., Drebot, M., Tyler, S., Shen, M., Cameron, C., Stoltz, D., et al. (2004). Complete nucleotide sequence of Kashmir bee virus and comparison with acute bee paralysis virus. Journal of General Virology, 85(8), 2263–2270.

Deleris, A., Gallego-Bartolome, J., Bao, J., Kasschau, K. D., Carrington, J. C., & Voinnet, O. (2006). Hierarchical action and inhibition of plant dicer-like proteins in antiviral defense. Science, 313(5783), 68–71.

Dhaygude, K., Johansson, H., Kulmuni, J., & Sundström, L. (2019). Genome organization and molecular characterization of the three Formica exsecta viruses—FeV1, FeV2 and FeV4. PeerJ, 6, e6216.

Edgar, R. C., Taylor, J., Lin, V., Altman, T., Barbera, P., Meleshko, D., et al. (2022). Petabase-scale sequence alignment catalyses viral discovery. Nature, 602(7895), 142–147.

Erickson, J. M. (1971). The displacement of native ant species by the introduced Argentine ant Iridomyrmex humilis Mayr. Psyche: A Journal of Entomology, 78(4), 257–266.

Evans, J. D., & Spivak, M. (2010). Socialized medicine: Individual and communal disease barriers in honey bees. Journal of Invertebrate Pathology, 103, S62–S72.

French, R. K., & Holmes, E. C. (2020). An Ecosystems Perspective on Virus Evolution and Emergence. Trends in Microbiology, 28, 165–175.

Fukasawa, F., Hirai, M., Takaki, Y., Shimane, Y., Thomas, C. E., Urayama, S., … Koyama, S. (2020). A new polycipivirus identified in Colobopsis shohki. Archives of Virology, 165(3), 761–763.

Fürst, M., McMahon, D. P., Osborne, J., Paxton, R., & Brown, M. (2014). Disease associations between honeybees and bumblebees as a threat to wild pollinators. Nature, 506(7488), 364.

Gammon, D. B., & Mello, C. C. (2015). RNA interference-mediated antiviral defense in insects. Current Opinion in Insect Science, 8, 111–120.

Geffre, A. C., Gernat, T., Harwood, G. P., Jones, B. M., Morselli Gysi, D., Hamilton, A. R., et al. (2020). Honey bee virus causes context-dependent changes in host social behavior. Proceedings of the National Academy of Sciences of the United States of America, 117(19), 10406–10413.

Giraud, T., Pedersen, J. S., & Keller, L. (2002). Evolution of supercolonies: The Argentine ants of southern Europe. Proceedings of the National Academy of Sciences of the United States of America, 99(9), 6075–6079.

Göertz, G. P., Miesen, P., Overheul, G. J., van Rij, R. P., van Oers, M. M., & Pijlman, G. P. (2019). Mosquito small RNA responses to West Nile and insect-specific virus infections in Aedes and Culex mosquito cells. Viruses, 11(3), 271.

Grabherr, M. G., Haas, B. J., Yassour, M., Levin, J. Z., Thompson, D. A., Amit, I., et al. (2011). Full-length transcriptome assembly from RNA-seq data without a reference genome. Nature Biotechnology, 29(7), 644–652.

Groden, E., Drummond, F. A., Garnas, J., & Franceour, A. (2005). Distribution of an invasive ant, Myrmica rubra (Hymenoptera: Formicidae), in Maine. Journal of Economic Entomology, 98(6), 1774–1784.

Grozinger, C. M., & Flenniken, M. L. (2019). Bee viruses: Ecology, pathogenicity, and impacts. Annual Review of Entomology, 64, 205–226.

Gruber, M. A. M., Cooling, M., Baty, J. W., Buckley, K., Friedlander, A., Quinn, O., et al. (2017). Single-stranded RNA viruses infecting the invasive Argentine ant, Linepithema humile. Scientific Reports, 7(1), 3304.

Guindon, S., Dufayard, J., Lefort, V., Anisimova, M., Hordijk, W., & Gascuel, O. (2010). New algorithms and methods to estimate maximum-likelihood phylogenies: Assessing the performance of PhyML 3.0. Systematic Biology, 59(3), 307–321.

Han, Y. H., Luo, Y. J., Wu, Q., Jovel, J., Wang, X. H., Aliyari, R., et al. (2011). RNA-based immunity terminates viral infection in adult Drosophila in the absence of viral suppression of RNA interference: Characterization of viral small interfering RNA populations in wild-type and mutant flies. Journal of Virology, 85(24), 13153–13163.

Hashimoto, Y., & Valles, S. M. (2008). Detection and quantitation of Solenopsis invicta virus-2 genomic and intermediary replicating viral RNA in fire ant workers and larvae. Journal of Invertebrate Pathology, 98(2), 243–245.

Hirakata, S., & Siomi, M. C. (2016). piRNA biogenesis in the germline: From transcription of piRNA genomic sources to piRNA maturation. Biochimica et Biophysica Acta, 1859(1), 82–92.

Holway, D. A., & Suarez, A. V. (1999). Animal behavior: An essential component of invasion biology. Trends in Ecology & Evolution, 14(8), 328–330.

Huang, X., & Madan, A. (1999). CAP3: A DNA sequence assembly program. Genome Research, 9(9), 868–877.

Hughes, W. O. H., Eilenberg, J., & Boomsma, J. J. (2002). Trade-offs in group living: Transmission and disease resistance in leaf-cutting ants. Proceedings of the Royal Society of London Series B-Biological Sciences, 269(1502), 1811–1819.

Hunt, M., Gall, A., Ong, S. H., Brener, J., Ferns, B., Goulder, P., et al. (2015). IVA: Accurate de novo assembly of RNA virus genomes. Bioinformatics, 31(14), 2374–2376.

Katoh, K., & Standley, D. M. (2013). MAFFT multiple sequence alignment software version 7: Improvements in performance and usability. Molecular Biology and Evolution, 30(4), 772–780.

Kemp, C., & Imler, J. (2009). Antiviral immunity in Drosophila. Current Opinion in Immunology, 21(1), 3–9.

Konrad, M., Vyleta, M. L., Theis, F. J., Stock, M., Tragust, S., Klatt, M., … Cremer, S. (2012). Social transfer of pathogenic fungus promotes active immunisation in ant colonies. PLoS Biology, 10(4), e1001300.

Langmead, B., & Salzberg, S. L. (2012). Fast gapped-read alignment with bowtie 2. Nature Methods, 9(4), 357–359.

Langmead, B., Trapnell, C., Pop, M., & Salzberg, S. L. (2009). Ultrafast and memory-efficient alignment of short DNA sequences to the human genome. Genome Biology, 10(3), R25.

Lee, Y. S., Nakahara, K., Pham, J. W., Kim, K., He, Z., Sontheimer, E. J., et al. (2004). Distinct roles for Drosophila dicer-1 and dicer-2 in the siRNA/miRNA silencing pathways. Cell, 117(1), 69–81.

Levitt, A. L., Singh, R., Cox-Foster, D. L., Rajotte, E., Hoover, K., Ostiguy, N., et al. (2013). Cross-species transmission of honey bee viruses in associated arthropods. Virus Research, 176(1-2), 232–240.

Li, H., & Durbin, R. (2009). Fast and accurate short read alignment with burrows-wheeler transform. Bioinformatics, 25(14), 1754–1760.

Li, H., Li, W. X., & Ding, S. W. (2002). Induction and suppression of RNA silencing by an animal virus. Science, 296(5571), 1319–1321.

Li, C.-X., Shi, M., Tian, J.-H., Lin, X.-D., Kang, Y.-J., Chen, L.-J., … Zhang, Y.-Z. (2015) Unprecedented genomic diversity of RNA viruses in arthropods reveals the ancestry of negative-sense RNA viruses. eLife, 4, e05378.

Lopez, W., Page, A. M., Carlson, D. J., Ericson, B. L., Cserhati, M. F., Guda, C., et al. (2018). Analysis of immune-related genes during Nora virus infection of Drosophila melanogaster using next generation sequencing. AIMS Microbiology, 4(1), 123–139.

Manfredini, F., Shoemaker, D., & Grozinger, C. M. (2016). Dynamic changes in host–virus interactions associated with colony founding and social environment in fire ant queens (Solenopsis invicta). Ecology and Evolution, 6(1), 233–244.

Martin, J. M. & Brettell, L. E. (2019). Deformed Wing Virus in Honeybees and Other Insects. Annual Review of Virology 6, 49–69.

McMahon, D. P., Fürst, M. A., Caspar, J., Theodorou, P., Brown, M. J., & Paxton, R. J. (2015). A sting in the spit: Widespread cross-infection of multiple RNA viruses across wild and managed bees. Journal of Animal Ecology, 84(3), 615–624.

McMenamin, A., Daughenbaugh, K., Parekh, F., Pizzorno, M., & Flenniken, M. (2018). Honey bee and bumble bee antiviral defense. Viruses, 10(8), 395.

Medd, N. C., Fellous, S., Waldron, F. M., Xuéreb, A., Nakai, M., Cross, J. V., et al. (2018). The virome of Drosophila suzukii, an invasive pest of soft fruit. Virus Evolution, 4(1), vey009.

Nayak, A., Berry, B., Tassetto, M., Kunitomi, M., Acevedo, A., Deng, C., et al. (2010). Cricket paralysis virus antagonizes Argonaute 2 to modulate antiviral defense in Drosophila. Nature Structural & Molecular Biology, 17(5), 547.

NCBI Resource Coordinators. (2017). Database resources of the National Center for Biotechnology Information. Nucleic Acids Research, 45(Database issue), D12–D17.

Oi, D., Porter, S., & Valles, S. (2015). A review of the biological control of fire ants. Myrmecological News, 21, 101–116.

Oi, D., Valles, S., Porter, S., Cavanaugh, C., White, G., & Henke, J. (2019). Introduction of fire ant biological control agents into the Coachella valley of California. Florida Entomologist, 102(1), 284–286.

Olendraite, I., Lukhovitskaya, N. I., Porter, S. D., Valles, S. M., & Firth, A. E. (2017). Polycipiviridae: A proposed new family of polycistronic picorna-like RNA viruses. Journal of General Virology, 98(9), 2368–2378.

Pettersson, J. H., Shi, M., Bohlin, J., Eldholm, V., Brynildsrud, O. B., Paulsen, K. M., et al. (2017). Characterizing the virome of Ixodes ricinus ticks from northern Europe. Scientific Reports, 7(1), 10870.

Pull, C. D., Metzler, S., Naderlinger, E., & Cremer, S. (2018). Protection against the lethal side effects of social immunity in ants. Current Biology, 28(19), R1139–R1140.

Pull, C. D., Ugelvig, L. V., Wiesenhofer, F., Grasse, A. V., Tragust, S., Schmitt, T., et al. (2018). Destructive disinfection of infected brood prevents systemic disease spread in ant colonies. Elife, 7, e32073.

Remnant, E. J., Shi, M., Buchmann, G., Blacquiere, T., Holmes, E. C., Beekman, M., et al. (2017). A diverse range of novel RNA viruses in geographically distinct honey bee populations. Journal of Virology, 91(16), 10.1128/JVI.00158-17

Rosengaus, R. B., Maxmen, A. B., Coates, L. E., & Traniello, J. F. (1998). Disease resistance: A benefit of sociality in the dampwood termite Zootermopsis angusticollis (isoptera: Termopsidae). Behavioral Ecology and Sociobiology, 44(2), 125–134.

Rothenbuhler, W. C. (1964). Behavior genetics of nest cleaning in honey bees. iv. responses of F1 and backcross generations to disease-killed brood. American Zoologist, 4, 111–123.

Sabin, L. R., Zheng, Q., Thekkat, P., Yang, J., Hannon, G. J., Gregory, B. D., et al. (2013). Dicer-2 processes diverse viral RNA species. PloS One, 8(2), e55458.

Schläppi, D., Chejanovsky, N., Yañez, O., & Neumann, P. (2020). Foodborne transmission and clinical symptoms of honey bee viruses in ants Lasius spp. Viruses, 12(3), 321.

Schläppi, D., Lattrell, P., Yañez, O., Chejanovsky, N., & Neumann, P. (2019). Foodborne transmission of deformed wing virus to ants (Myrmica rubra). Insects, 10(11), 394.

Schmid-Hempel, P. (1998). Parasites in social insects. Princeton University Press.

Seifert, B. (2000). Rapid range expansion in Lasius neglectus (Hymenoptera, formicidae)—an Asian invader swamps Europe. Deutsche Entomologische Zeitschrift, 47(2), 173–179.

Shi, M., Lin, X., Tian, J., Chen, L., Chen, X., Li, C., et al. (2016). Redefining the invertebrate RNA virosphere. Nature, 540(7634), 539–543.

Singh, R., Levitt, A. L., Rajotte, E. G., Holmes, E. C., Ostiguy, N., Lipkin, W. I., et al. (2010). RNA viruses in hymenopteran pollinators: Evidence of inter-taxa virus transmission via pollen and potential impact on non-apis hymenopteran species. PloS One, 5(12), e14357.

Siva-Jothy, M. T., Moret, Y., & Rolff, J. (2005). Insect immunity: An evolutionary ecology perspective. Advances in Insect Physiology, 32, 1–48.

Son, K. N., Liang, Z., & Lipton, H. L. (2015). Double-stranded RNA is detected by immunofluorescence analysis in RNA and DNA virus infections, including those by negative-stranded RNA viruses. Journal of Virology, 89(18), 9383–9392.

Stroeymeyt, N., Grasse, A. V., Crespi, A., Mersch, D. P., Cremer, S., & Keller, L. (2018). Social network plasticity decreases disease transmission in a eusocial insect. Science, 362(6417), 941–945.

Suarez, A. V., Holway, D. A., & Case, T. J. (2001). Patterns of spread in biological invasions dominated by long-distance jump dispersal: Insights from Argentine ants. Proceedings of the National Academy of Sciences of the United States of America, 98(3), 1095–1100.

Theis, F. J., Ugelvig, L. V., Marr, C., & Cremer, S. (2015). Opposing effects of allogrooming on disease transmission in ant societies. Philosophical Transactions of the Royal Society of London. Series B, Biological Sciences, 370(1669), 10.1098/rstb.2014.0108.

Tragust, S., Ugelvig, L. V., Chapuisat, M., Heinze, J., & Cremer, S. (2013). Pupal cocoons affect sanitary brood care and limit fungal infections in ant colonies. BMC Evolutionary Biology, 13, 225.

Ugelvig, L. V., & Cremer, S. (2012). Effects of social immunity and unicoloniality on host– parasite interactions in invasive insect societies. Functional Ecology, 26(6), 1300–1312.

Ugelvig, L. V., Drijfhout, F. P., Kronauer, D. J., Boomsma, J. J., Pedersen, J. S., & Cremer, S. (2008). The introduction history of invasive garden ants in Europe: Integrating genetic, chemical and behavioural approaches. BMC Biology, 6(1), 11.

Valles, S. M., & Hashimoto, Y. (2009). Isolation and characterization of Solenopsis invicta virus 3, a new positive-strand RNA virus infecting the red imported fire ant, Solenopsis invicta. Virology, 388(2), 354–361.

Valles, S. M., Oi, D. H., Becnel, J. J., Wetterer, J. K., LaPolla, J. S., & Firth, A. E. (2016). Isolation and characterization of Nylanderia fulva virus 1, a positive-sense, single-stranded RNA virus infecting the tawny crazy ant, Nylanderia fulva. Virology, 496, 244–254.

Valles, S. M., Oi, D. H., Yu, F., Tan, X., & Buss, E. A. (2012). Metatranscriptomics and pyrosequencing facilitate discovery of potential viral natural enemies of the invasive caribbean crazy ant, Nylanderia pubens. PloS One, 7(2), e31828.

Valles, S. M., Porter, S. D., & Calcaterra, L. A. (2018). Prospecting for viral natural enemies of the fire ant Solenopsis invicta in Argentina. PloS One, 13(2), e0192377.

Valles, S. M., & Rivers, A. R. (2019). Nine new RNA viruses associated with the fire ant Solenopsis invicta from its native range. Virus Genes, 55(5), 368–380.

Valles, S. M., Strong, C. A., Dang, P. M., Hunter, W. B., Pereira, R. M., Oi, D. H., et al. (2004). A picorna-like virus from the red imported fire ant, Solenopsis invicta: Initial discovery, genome sequence, and characterization. Virology, 328(1), 151–157.

Valles, S. M., Strong, C. A., & Hashimoto, Y. (2007). A new positive-strand RNA virus with unique genome characteristics from the red imported fire ant, Solenopsis invicta. Virology, 365(2), 457–463.

van Mierlo, J. T., Bronkhorst, A. W., Overheul, G. J., Sadanandan, S. A., Ekstrom, J. O., Heestermans, M., et al. (2012). Convergent evolution of Argonaute-2 slicer antagonism in two distinct insect RNA viruses. PLoS Pathogens, 8(8), e1002872.

van Rij, R. P., Saleh, M. C., Berry, B., Foo, C., Houk, A., Antoniewski, C., et al. (2006). The RNA silencing endonuclease Argonaute 2 mediates specific antiviral immunity in Drosophila melanogaster. Genes & Development, 20(21), 2985–2995.

Viljakainen, L., Holmberg, I., Abril, S., & Jurvansuu, J. (2018). Viruses of invasive argentine ants from the European main supercolony: Characterization, interactions and evolution. Journal of General Virology, 99, 1129–1140.

Watson, M., Schnettler, E., & Kohl, A. (2013). viRome: An R package for the visualization and analysis of viral small RNA sequence datasets. Bioinformatics, 29(15), 1902–1903.

Webster, C. L., Waldron, F. M., Robertson, S., Crowson, D., Ferrari, G., Quintana, J. F., et al. (2015). The discovery, distribution, and evolution of viruses associated with Drosophila melanogaster. PLoS Biology, 13(7), e1002210.

Wilfert, L., Long, G., Leggett, H. C., Schmid-Hempel, P., Butlin, R., Martin, S. J. M. & Boots, M. (2016) Deformed wing virus is a recent global epidemic in honeybees driven by Varroa mites. Science, 351(6273): 594–597

Wilson, R. C., & Doudna, J. A. (2013). Molecular mechanisms of RNA interference. Annual Review of Biophysics, 42, 217–239.

Xavier, C. A. D., Allen, M. L., & Whitfield, A. E. (2021) Ever-increasing viral diversity associated with the red imported fire ant Solenopsis invicta (Formicidae: Hymenoptera). Virology Journal 18, 5.

Yilmaz, P., Parfrey, L. W., Yarza, P., Gerken, J., Pruesse, E., Quast, C., et al. (2013). The SILVA and “all-species living tree project (LTP)” taxonomic frameworks. Nucleic Acids Research, 42(D1), D643–D648.

Zell, R., Delwart, E., Gorbalenya, A. E., Hovi, T., King, A. M. Q., Knowles, N. J., et al. (2017). ICTV virus taxonomy profile: Picornaviridae. The Journal of General Virology, 98(10), 2421–2422.

Zheng, Y., Gao, S., Padmanabhan, C., Li, R., Galvez, M., Gutierrez, D., et al. (2017). VirusDetect: An automated pipeline for efficient virus discovery using deep sequencing of small RNAs. Virology, 500, 130–138.

