## Supplementary material for "Antiviral immune response reveals host-specific virus infections in natural ant populations"

**for**

This file contains

- supplemental tables 1-2 (table S3 separately in xlsx format)
- supplemental figures 1-4

**Supplemental Table 1. RNA viruses discovered in ants prior to this study.** List of viruses of ants published in 2004-2021. Columns provide information on the host ant species, the published virus name, and its phylogenetic classification (to the known level of order, family or genus). The last column gives reference to the publication that described the virus.

| Host ant | Virus name | Classification[a] | Publication |
|---|---|---|---|
| *Camponotus japonicus* | *Wuhan ant virus* | *Mononegavirales; Rhabdoviridae* | Li et al., 2015 |
| *Colobopsis shohki* | *Colobopsis shohki virus 1* | *Picornavirales; Polycipiviridae; Sopolycivirus* | Fukasawa et al., 2020 |
| *Formica fusca* | *Formica fusca virus 1* | *Nyamiviridae* | Kleanthous, Olendraite, Lukhovitskaya, & Firth, 2019 |
| *F. exsecta* | *Formica exsecta virus-1* | *Picornavirales; Dicistroviridae* | Dhaygude, Johansson, Kulmuni, & Sundström, 2019 |
| *F. exsecta* | *Formica exsecta virus-2* | *Picornavirales; Iflaviridae* | Dhaygude et al., 2019 |
| *F. exsecta* | *Formica exsecta* TSA | *Picornavirales; Polycipiviridae; Sopolycivirus* | Olendraite, Lukhovitskaya, Porter, Valles, & Firth, 2017 |
| *F. exsecta* | *Formica exsecta virus-4* | *Mononegavirales* | Dhaygude et al., 2019 |
| *Linepithema humile* | *Kashmir bee virus* | *Picornavirales; Dicistroviridae; Aparavirus* | Gruber et al., 2017; Viljakainen, Holmberg, Abril, & Jurvansuu, 2018 |
| *Li. humile* | *Linepithema humile bunya-like virus 1* | *Bunyavirales* | Viljakainen et al., 2018 |
| *Li. humile* | *Linepithema humile C-virus 1* | Unclassified | Viljakainen et al., 2018 |
| *Li. humile* | *Linepithema humile narna-like virus 1* | *Narnaviridae* | Viljakainen et al., 2018 |
| *Li. humile* | *Linepithema humile polycipivirus 1* | *Picornavirales; Polycipiviridae; Sopolycivirus* | Olendraite et al., 2017; Viljakainen et al., 2018 |
| *Li. humile* | *Linepithema humile polycipivirus 2* | *Picornavirales; Polycipiviridae; Sopolycivirus* | Viljakainen et al., 2018 |
| *Li. humile* | *Linepithema humile picorna-like virus 1* | *Picornavirales* | Viljakainen et al., 2018 |
| *Li. humile* | *Linepithema humile partiti-like virus 1* | *Mononegavirales; Partitiviridae* | Viljakainen et al., 2018 |
| *Li. humile* | *Linepithema humile rhabdo-like virus 1* | *Mononegavirales; Rhabdoviridae* | Viljakainen et al., 2018 |
| *Li. humile* | *Linepithema humile toti-like virus 1* | *Totiviridae* | Viljakainen et al., 2018 |
| *Li. humile* | *Linepithema humile virus 1* | *Picornavirales* | Gruber et al., 2017; Viljakainen et al., 2018 |
| *Lasius neglectus* | *Lasius neglectus virus 1* | *Picornavirales; Polycipiviridae; Sopolycivirus* | Olendraite et al., 2017 |
| *La. neglectus* | *Lasius neglectus* TSA | *Picornavirales; Polycipiviridae; Sopolycivirus* | Olendraite et al., 2017 |
| *La. neglectus* | *Lasius neglectus virus 2* | *Mononegavirales; Rhabdoviridae* | Kleanthous et al., 2019 |
| *La. niger* | *Lasius niger virus 1* | *Picornavirales; Polycipiviridae; Sopolycivirus* | Olendraite et al., 2017 |
| *Monomorium pharaonis* | *Monomorium pharaonis* TSA | *Picornavirales; Polycipiviridae; Sopolycivirus* | Olendraite et al., 2017 |
| *Myrmica scabrinodis* | *Myrmica scabrinodis virus 1* | *Picornavirales; Polycipiviridae; Sopolycivirus* | Olendraite et al., 2017 |
| *M. scabrinodis* | *Myrmica scabrinodis virus 2* | *Picornavirales; Dicistroviridae* | Kleanthous et al., 2019 |
| *Nylanderia fulva* | *Nylanderia fulva virus 1* | *Solinviviridae; Nyfulvavirus* | Valles et al., 2016 |
| *N. pubens* | - | Three unnamed viruses | Valles, Oi, Yu, Tan, & Buss, 2012 |
| *Solenopsis invicta* | *Solenopsis invicta virus 1* | *Picornavirales; Dicistroviridae; Aparavirus* | Valles et al., 2004 |
| *S. invicta* | *Solenopsis invicta virus 2* | *Picornavirales; Polycipiviridae; Sopolycivirus* | Olendraite et al., 2017; Valles, Strong, & Hashimoto, 2007 |
| *S. invicta* | *Solenopsis invicta virus 3* | *Solinviviridae* | Valles & Hashimoto, 2009 |
| *S. invicta* | *Solenopsis invicta virus 4* | *Picornavirales; Polycipiviridae; Sopolycivirus* | Olendraite et al., 2017 |
| *S. invicta* | *Solenopsis invicta virus 5* | *Picornavirales; Dicistroviridae* | Valles, Porter, & Calcaterra, 2018 |
| *S. invicta* | *Solenopsis invicta virus 6* | *Picornavirales; Dicistroviridae; Cripavirus* | Valles & Rivers, 2019 |
| *S. invicta* | *Solenopsis invicta virus 7* | Unclassified | Valles & Rivers, 2019 |
| *S. invicta* | *Solenopsis invicta virus 8* | *Picornavirales; Polycipiviridae; Sopolycivirus* | Valles & Rivers, 2019 |
| *S. invicta* | *Solenopsis invicta virus 9* | *Picornavirales; Dicistroviridae; Triatovirus* | Valles & Rivers, 2019 |
| *S. invicta* | *Solenopsis invicta virus 10* | Unclassified | Valles & Rivers, 2019 |
| *S. invicta* | *Solenopsis invicta virus 11* | *Picornavirales; Iflaviridae* | Valles & Rivers, 2019 |
| *S. invicta* | *Solenopsis invicta virus 12* | *Picornavirales; Dicistroviridae; Aparavirus* | Valles & Rivers, 2019 |
| *S. invicta* | *Solenopsis invicta virus 13* | *Picornavirales; Dicistroviridae; Aparavirus* | Valles & Rivers, 2019 |
| *S. invicta* | *Solenopsis invicta virus 14* | *Bunyavirales* | Xavier, Allen, & Whitfield, 2021 |
| *S. invicta* | *Solenopsis invicta virus 15* | *Mononegavirales* | Xavier, Allen, & Whitfield, 2021 |
| *S. invicta* | *Solenopsis invicta virus 16* | *Picornavirales; Iflaviridae* | Xavier, Allen, & Whitfield, 2021 |
| *S. invicta* | *Solenopsis invicta virus 17* | Unclassified | Xavier, Allen, & Whitfield, 2021 |
| *S. invicta* | *Solenopsis midden virus* | *Totiviridae* | Valles & Rivers, 2019 |
| *Tapinoma melanocephalum* | *Milolii virus* | Unclassified | Brettell, Mordecai, Pachori, & Martin, 2017 |

[a] presented in the order, family or genus level.

**Supplemental Table 2. Collection sites and sample composition of the ants**. For each population of the three ant species, the collection year, country, village/city and coordinates of the collection site are given. In addition, the sample composition, i.e. the number or workers, queens and brood items, that were pooled to obtain the sample, are provided.

| Ant species | population | year | country | village/city | coordinates | Workers | Queens | Brood |
|---|---|---|---|---|---|---|---|---|
| *Linepithema humile* | 1 | 2014 | Italy | Orbetello | 42°26'33.2"N 11°13'39.9"E | 500 | | |
| *Linepithema humile* | 2 | 2014 | Spain | L'Escala | 42°07'18.1"N 3°08'17.0"E | 550 | | |
| *Linepithema humile* | 3 | 2014 | Spain | Sant Feliu de Guíxols | 41°48'54.8"N 3°03'10.5"E | 500 | | 50 |
| *Lasius neglectus* | 1 | 2014 | Italy | Volterra | 43°24'00.0"N 10°51'00.0"E | 405 | | 107 |
| *Lasius neglectus* | 2 | 2014 | Spain | L'Escala | 42°07'18.1"N 3°08'17.0"E | 550 | | |
| *Lasius neglectus* | 3 | 2014 | Spain | Seva | 41°48'33.1"N 2°15'46.9"E | 500 | | 56 |
| *Myrmica rubra* | 1 | 2014 | Italy | Monza | 45°35'54.5"N 9°16'18.5"E | 433 | 1 | 75 |
| *Myrmica rubra* | 2 | 2014 | Spain | Ripoll | 42°12' 3"N 2°11'26"E | 527 | | 22 |
| *Myrmica rubra* | 3 | 2014 | Spain | Vilallonga de Ter | 42°19'43.3"N 2°18'41.5"E | 500 | | 65 |

**Supplemental Table 3. Virus abundance and host response.** For each population of each of the three ant species, we provide details (sequence length, # mapped reads, RPKM, depth, fraction of viral reads) of the viral abundances (based on RNA-seq) and the triggered RNAi host response (based on sRNA-seq) for all actively infecting viruses (virus names in bold) and for potential contaminant viruses.

*see Viljakainen_TableS3.xlsx*

**Supplemental Figure 1**

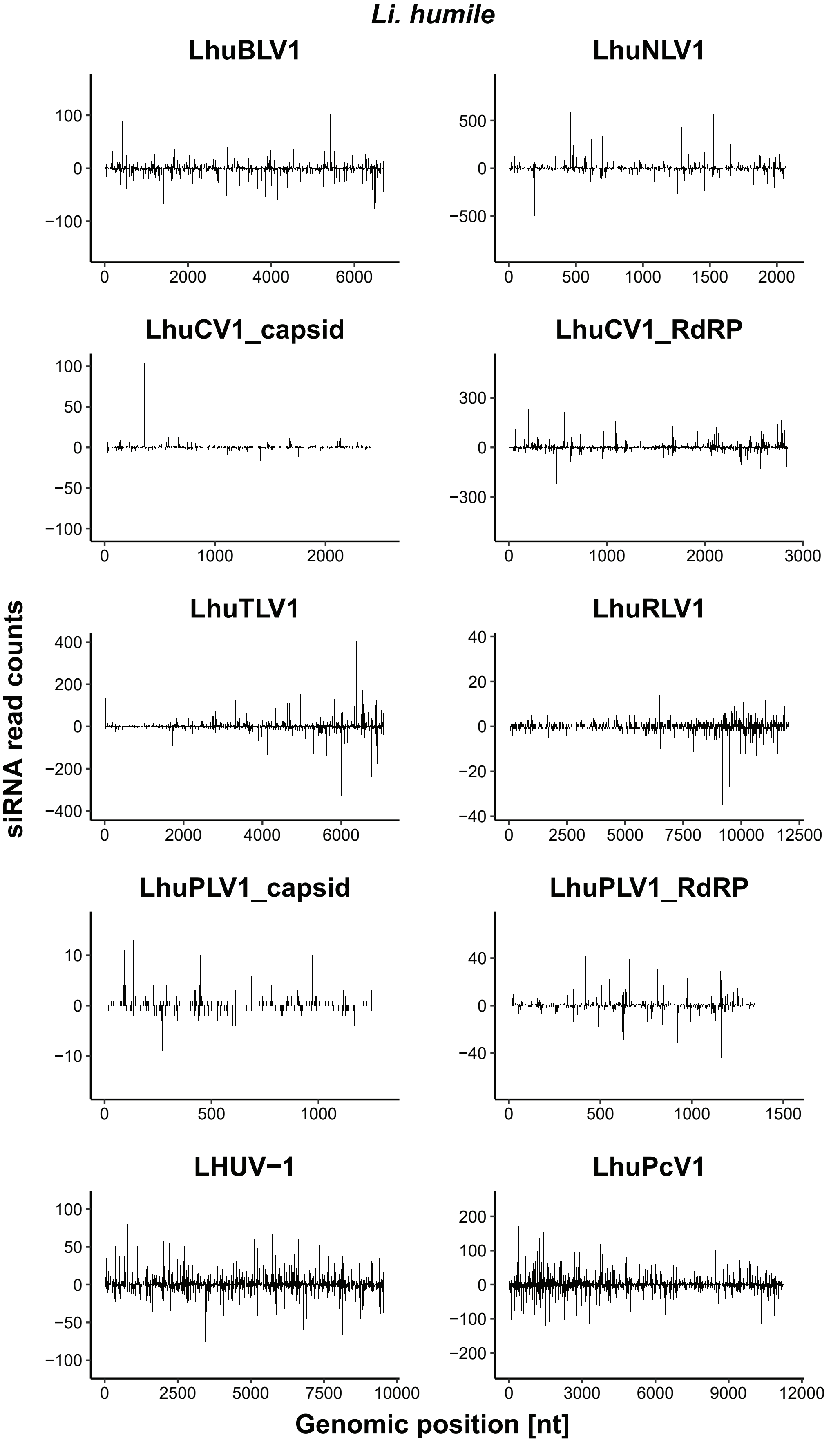

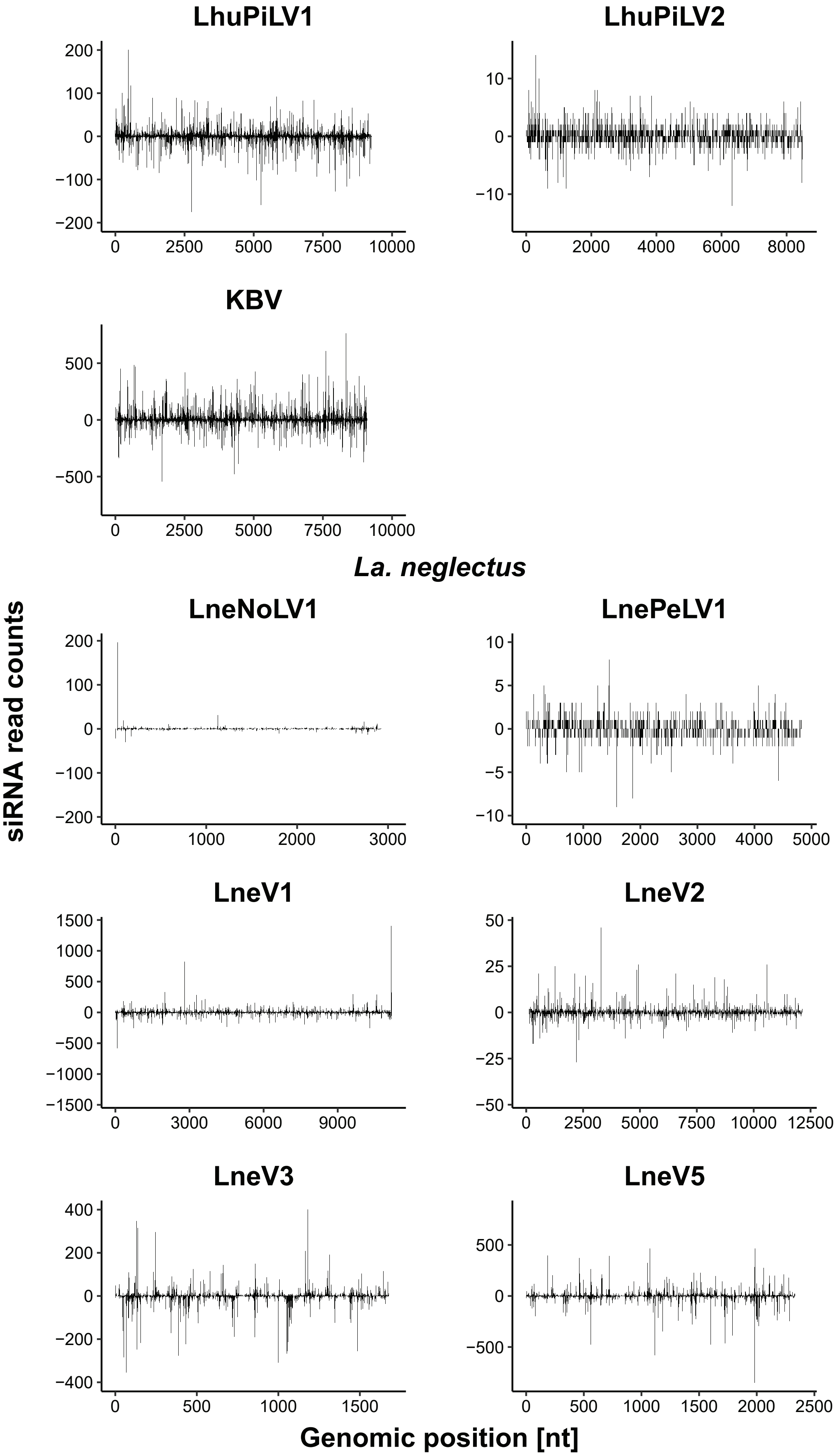
LhuPiLV1
LhuPiLV2
KBV
La. neglectus
LneNoLV1
LnePeLV1
LneV1
LneV2
LneV3
LneV5
siRNA read counts
Genomic position [nt]

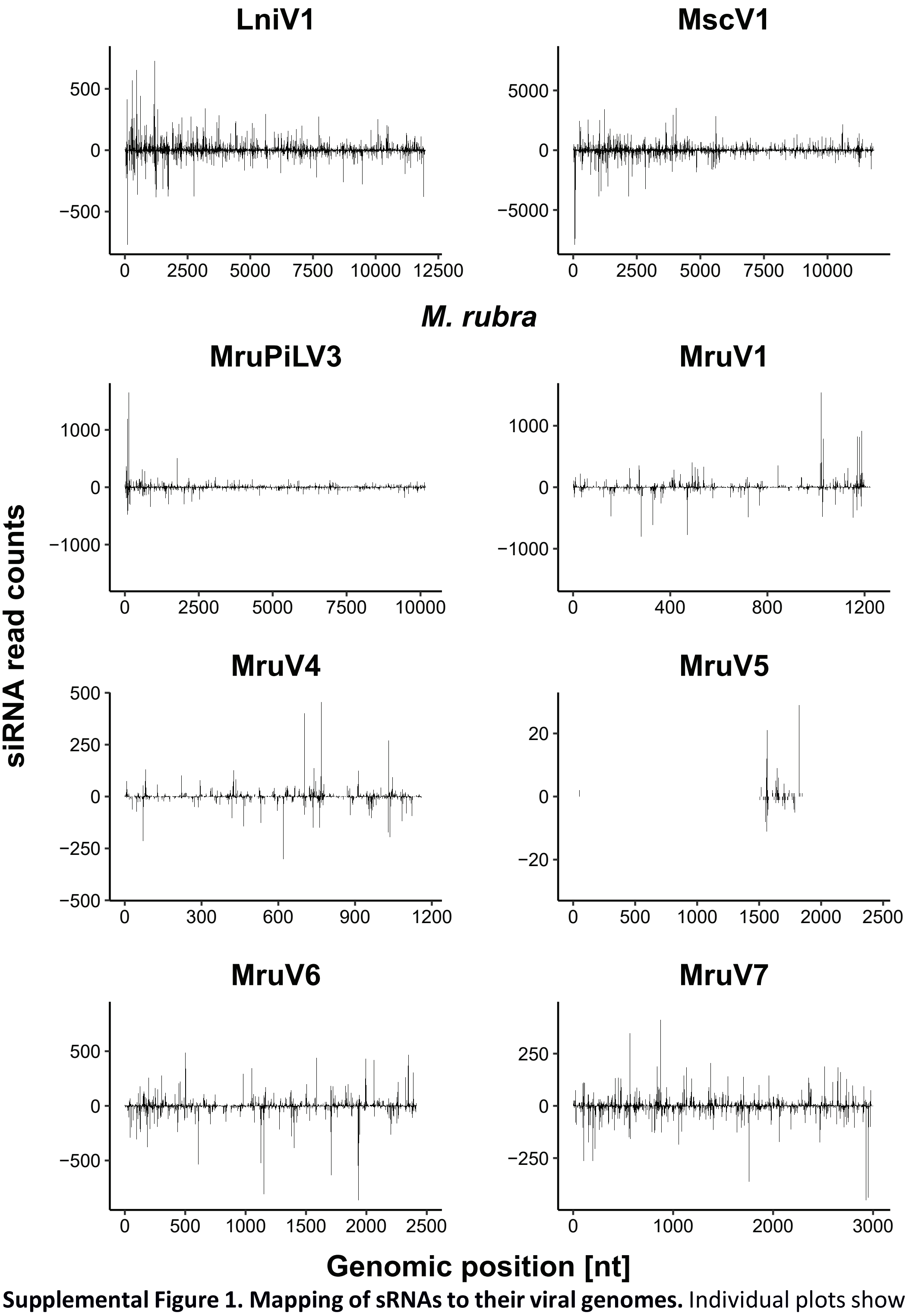


**Supplemental Figure 1. Mapping of sRNAs to their viral genomes.** Individual plots show the mapping of the small RNAs (sRNAs with confirmed proportion of 21-22nt being ≥ 50%) to their viral genomes assembled from the long RNA-seq data of the respective virus found in A) *Li. humile*, B) *La. neglectus* and C) *M. rubra* ants. All sequences are based on the complete virus genome assembly (two virus genomes, LhuCV1 and LhuPLV1 are bipartite). In total, 25 of the viruses (14 known and 11 novel) elicited a detectable RNAi response. For each virus, the x-axis shows the genomic position and y-axis the count of reads that map to each position. Reads that map to the positive strand are plotted above the x-axis and reads that map to the negative strand below the x-axis.

**Supplemental Figure 2**

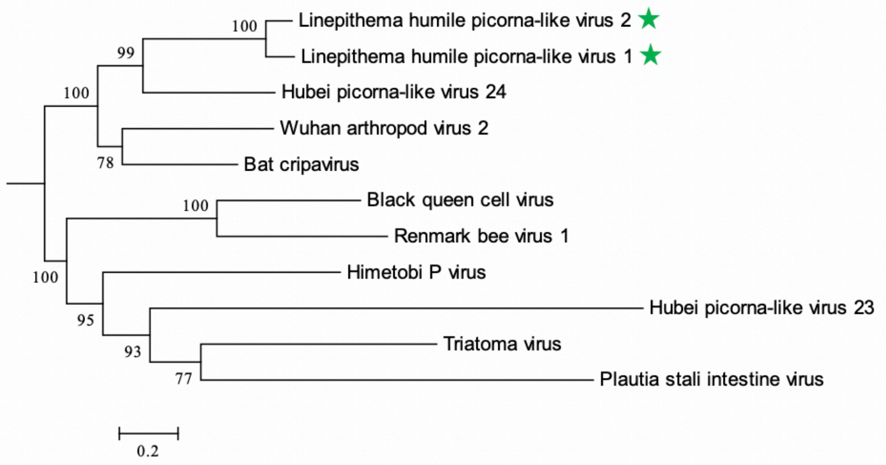
Linepithema humile picorna-like virus 2
Linepithema humile picorna-like virus 1
Hubei picorna-like virus 24
Wuhan arthropod virus 2
Bat cripavirus
Black queen cell virus
Renmark bee virus 1
Himetobi P virus
Hubei picorna-like virus 23
Triatoma virus
Plautia stali intestine virus
0.2

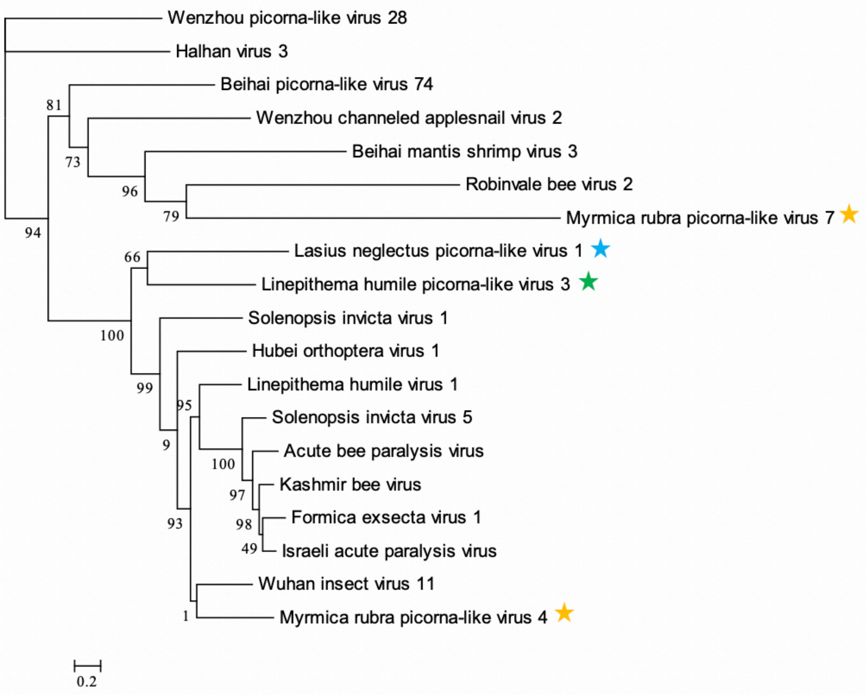
Wenzhou picorna-like virus 28
Halhan virus 3
Beihai picorna-like virus 74
Wenzhou channeled applesnail virus 2
Beihai mantis shrimp virus 3
Robinvale bee virus 2
Myrmica rubra picorna-like virus 7
Lasius neglectus picorna-like virus 1
Linepithema humile picorna-like virus 3
Solenopsis invicta virus 1
Hubei orthoptera virus 1
Linepithema humile virus 1
Solenopsis invicta virus 5
Acute bee paralysis virus
Kashmir bee virus
Formica exsecta virus 1
Israeli acute paralysis virus
Wuhan insect virus 11
Myrmica rubra picorna-like virus 4
0.2

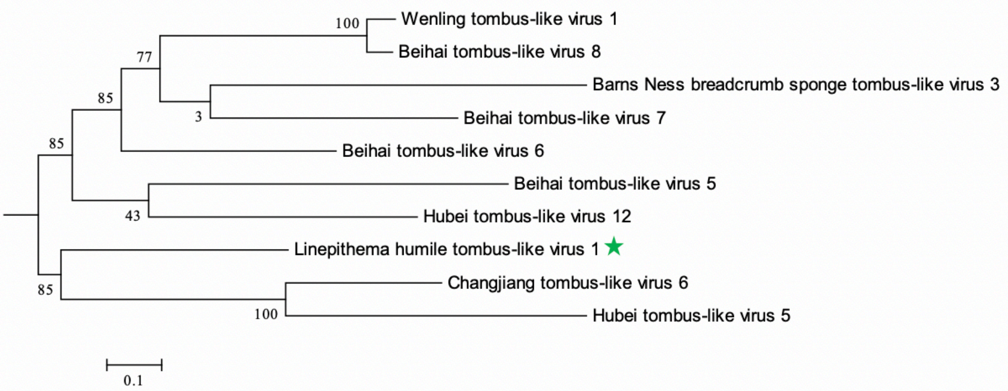
Wenling tombus-like virus 1
Beihai tombus-like virus 8
Barns Ness breadcrumb sponge tombus-like virus 3
Beihai tombus-like virus 7
Beihai tombus-like virus 6
Beihai tombus-like virus 5
Hubei tombus-like virus 12
Linepithema humile tombus-like virus 1
Changjiang tombus-like virus 6
Hubei tombus-like virus 5
0.1

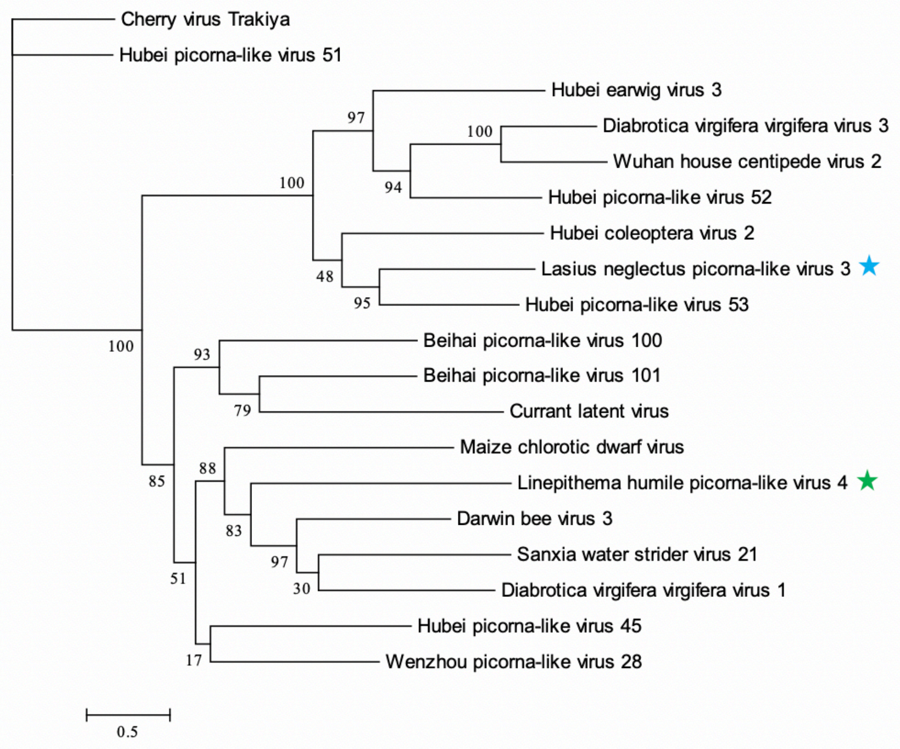
Cherry virus Trakiya
Hubei picorna-like virus 51
Hubei earwig virus 3
Diabrotica virgifera virgifera virus 3
Wuhan house centipede virus 2
Hubei picorna-like virus 52
Hubei coleoptera virus 2
Lasius neglectus picorna-like virus 3
Hubei picorna-like virus 53
Beihai picorna-like virus 100
Beihai picorna-like virus 101
Currant latent virus
Maize chlorotic dwarf virus
Linepithema humile picorna-like virus 4
Darwin bee virus 3
Sanxia water strider virus 21
Diabrotica virgifera virgifera virus 1
Hubei picorna-like virus 45
Wenzhou picorna-like virus 28
97
100
100
94
48
95
100
93
79
88
85
83
97
30
51
17
0.5

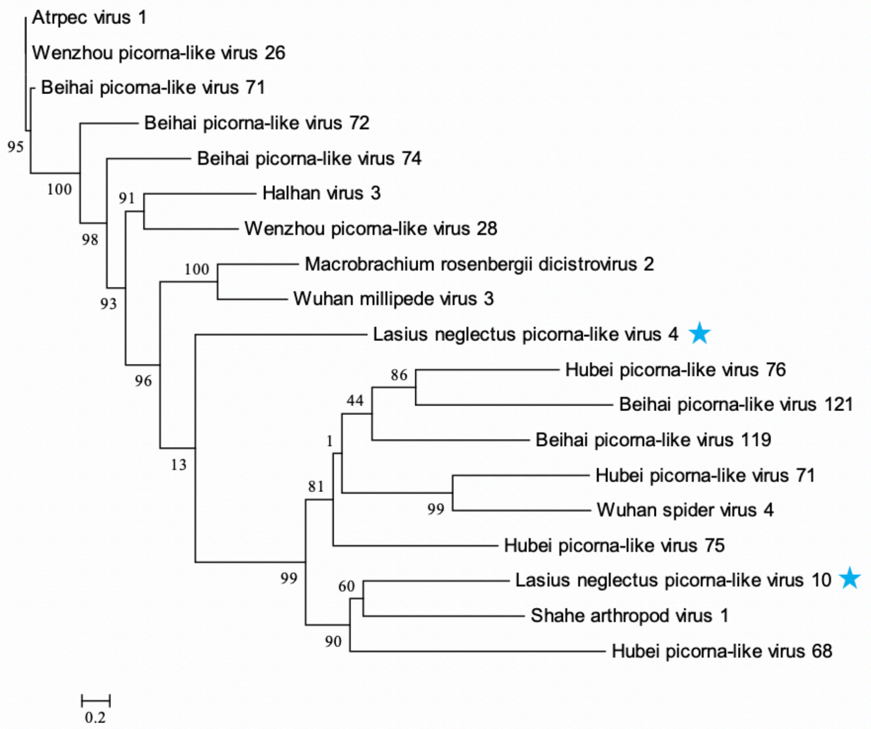
Atrpec virus 1
Wenzhou picorna-like virus 26
Beihai picorna-like virus 71
Beihai picorna-like virus 72
Beihai picorna-like virus 74
Halhan virus 3
Wenzhou picorna-like virus 28
Macrobrachium rosenbergii dicistrovirus 2
Wuhan millipede virus 3
Lasius neglectus picorna-like virus 4
Hubei picorna-like virus 76
Beihai picorna-like virus 121
Beihai picorna-like virus 119
Hubei picorna-like virus 71
Wuhan spider virus 4
Hubei picorna-like virus 75
Lasius neglectus picorna-like virus 10
Shahe arthropod virus 1
Hubei picorna-like virus 68
95
100
98
91
93
100
96
13
86
44
1
81
99
99
60
90
0.2

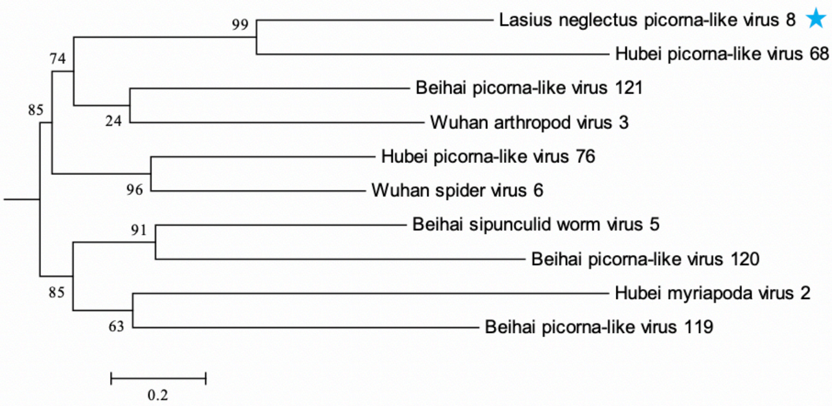

Lasius neglectus picorna-like virus 8
Hubei picorna-like virus 68
Beihai picorna-like virus 121
Wuhan arthropod virus 3
Hubei picorna-like virus 76
Wuhan spider virus 6
Beihai sipunculid worm virus 5
Beihai picorna-like virus 120
Hubei myriapoda virus 2
Beihai picorna-like virus 119
99
74
85
24
96
91
85
63
0.2


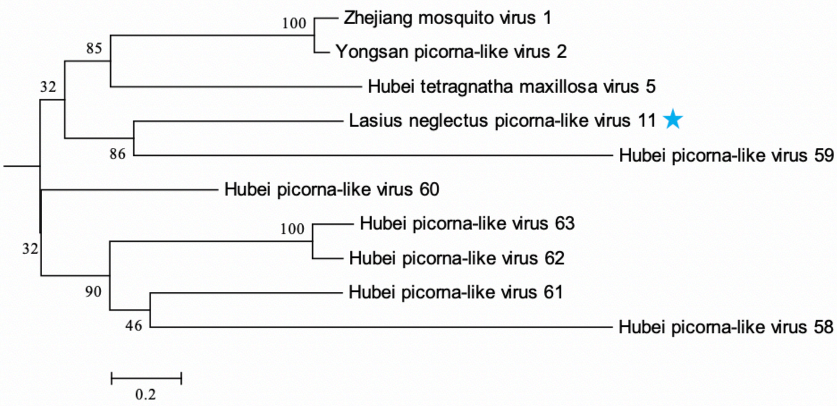

Zhejiang mosquito virus 1
Yongsan picorna-like virus 2
Hubei tetragnatha maxillosa virus 5
Lasius neglectus picorna-like virus 11
Hubei picorna-like virus 59
Hubei picorna-like virus 60
Hubei picorna-like virus 63
Hubei picorna-like virus 62
Hubei picorna-like virus 61
Hubei picorna-like virus 58
100
85
32
86
32
100
90
46
0.2


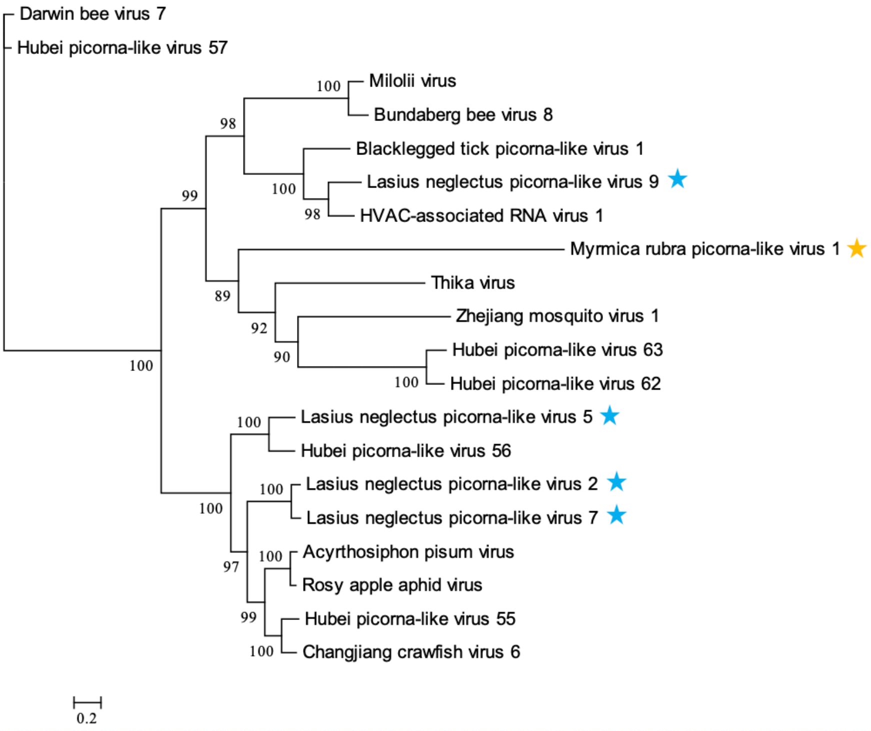

Darwin bee virus 7
Hubei picorna-like virus 57
Milolii virus
Bundaberg bee virus 8
Blacklegged tick picorna-like virus 1
Lasius neglectus picorna-like virus 9
HVAC-associated RNA virus 1
Myrmica rubra picorna-like virus 1
Thika virus
Zhejiang mosquito virus 1
Hubei picorna-like virus 63
Hubei picorna-like virus 62
Lasius neglectus picorna-like virus 5
Hubei picorna-like virus 56
Lasius neglectus picorna-like virus 2
Lasius neglectus picorna-like virus 7
Acyrthosiphon pisum virus
Rosy apple aphid virus
Hubei picorna-like virus 55
Changjiang crawfish virus 6
100
98
99
100
98
89
92
90
100
100
100
100
100
97
100
99
100
0.2

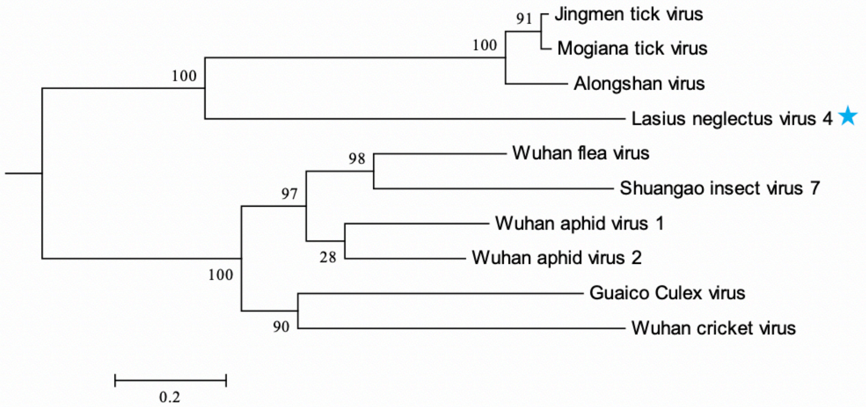
Jingmen tick virus
Mogiana tick virus
Alongshan virus
Lasius neglectus virus 4
Wuhan flea virus
Shuangao insect virus 7
Wuhan aphid virus 1
Wuhan aphid virus 2
Guaico Culex virus
Wuhan cricket virus
0.2

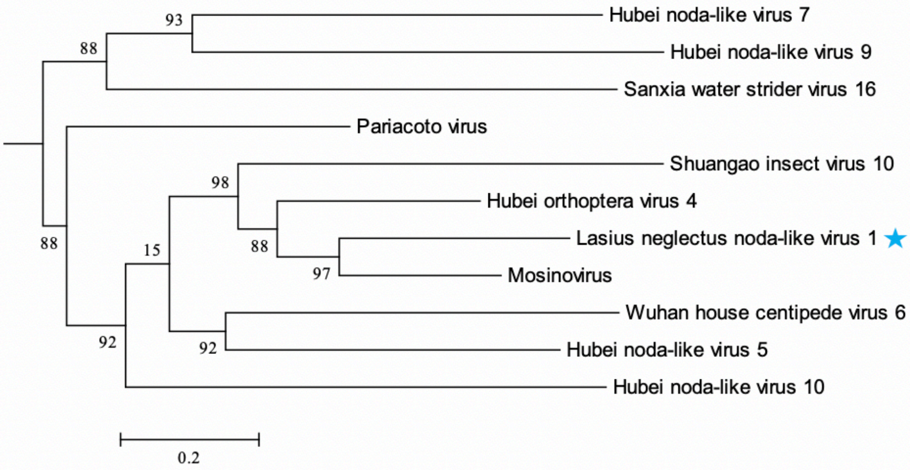
Hubei noda-like virus 7
Hubei noda-like virus 9
Sanxia water strider virus 16
Pariacoto virus
Shuangao insect virus 10
Hubei orthoptera virus 4
Lasius neglectus noda-like virus 1
Mosinovirus
Wuhan house centipede virus 6
Hubei noda-like virus 5
Hubei noda-like virus 10
0.2

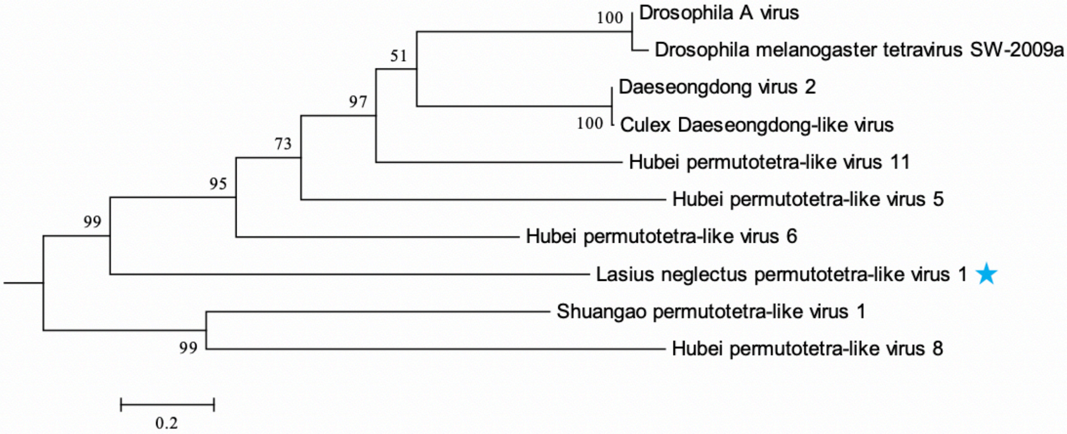
Drosophila A virus
Drosophila melanogaster tetravirus SW-2009a
Daeseongdong virus 2
Culex Daeseongdong-like virus
Hubei permutotetra-like virus 11
Hubei permutotetra-like virus 5
Hubei permutotetra-like virus 6
Lasius neglectus permutotetra-like virus 1
Shuangao permutotetra-like virus 1
Hubei permutotetra-like virus 8
0.2

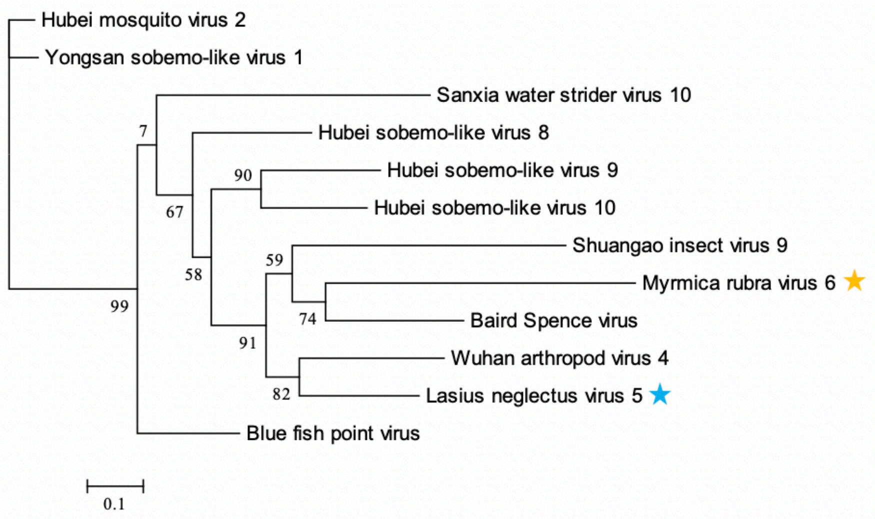
Hubei mosquito virus 2
Yongsan sobemo-like virus 1
Sanxia water strider virus 10
Hubei sobemo-like virus 8
Hubei sobemo-like virus 9
Hubei sobemo-like virus 10
Shuangao insect virus 9
Myrmica rubra virus 6
Baird Spence virus
Wuhan arthropod virus 4
Lasius neglectus virus 5
Blue fish point virus
7
90
67
59
58
74
99
91
82
0.1

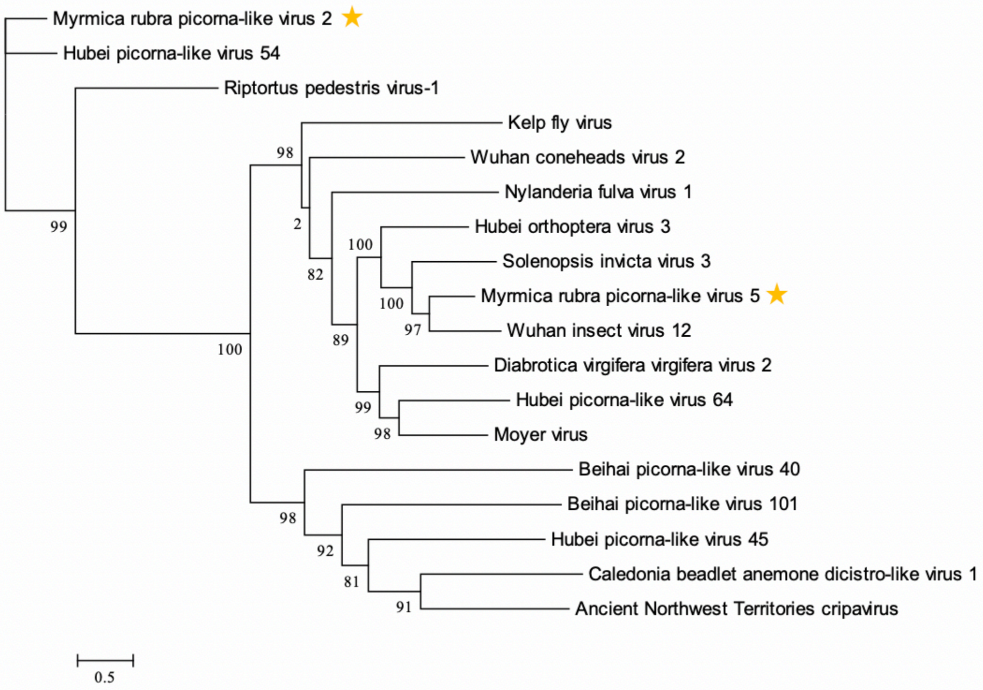
Myrmica rubra picorna-like virus 2
Hubei picorna-like virus 54
Riptortus pedestris virus-1
Kelp fly virus
Wuhan coneheads virus 2
Nylanderia fulva virus 1
Hubei orthoptera virus 3
Solenopsis invicta virus 3
Myrmica rubra picorna-like virus 5
Wuhan insect virus 12
Diabrotica virgifera virgifera virus 2
Hubei picorna-like virus 64
Moyer virus
Beihai picorna-like virus 40
Beihai picorna-like virus 101
Hubei picorna-like virus 45
Caledonia beadlet anemone dicistro-like virus 1
Ancient Northwest Territories cripavirus
98
2
100
82
100
97
89
99
98
99
100
98
92
81
91
0.5

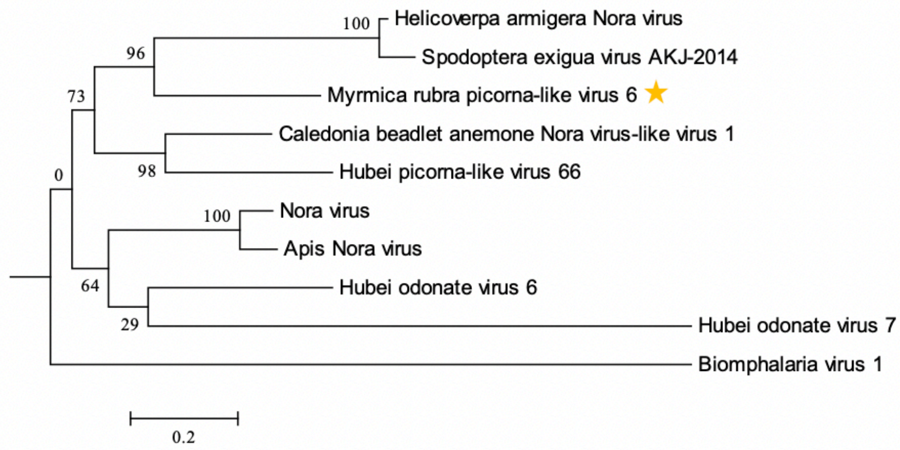
100
Helicoverpa armigera Nora virus
96
Spodoptera exigua virus AKJ-2014
73
Myrmica rubra picorna-like virus 6
Caledonia beadlet anemone Nora virus-like virus 1
0
98
Hubei picorna-like virus 66
100
Nora virus
Apis Nora virus
64
Hubei odonate virus 6
29
Hubei odonate virus 7
Biomphalaria virus 1
0.2

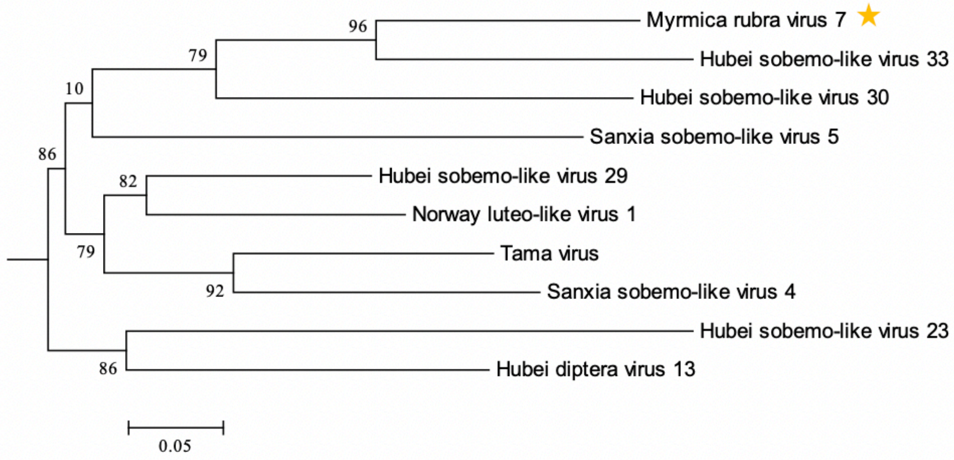
96
Myrmica rubra virus 7
79
Hubei sobemo-like virus 33
10
Hubei sobemo-like virus 30
Sanxia sobemo-like virus 5
86
82
Hubei sobemo-like virus 29
Norway luteo-like virus 1
79
Tama virus
92
Sanxia sobemo-like virus 4
Hubei sobemo-like virus 23
86
Hubei diptera virus 13
0.05

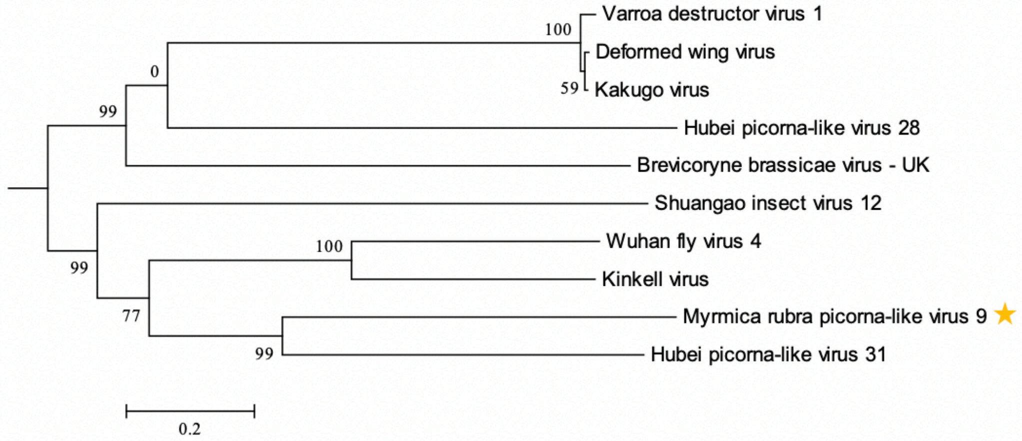
Varroa destructor virus 1
100
Deformed wing virus
0
59
Kakugo virus
99
Hubei picorna-like virus 28
Brevicoryne brassicae virus - UK
Shuangao insect virus 12
100
Wuhan fly virus 4
99
Kinkell virus
77
Myrmica rubra picorna-like virus 9
99
Hubei picorna-like virus 31
0.2

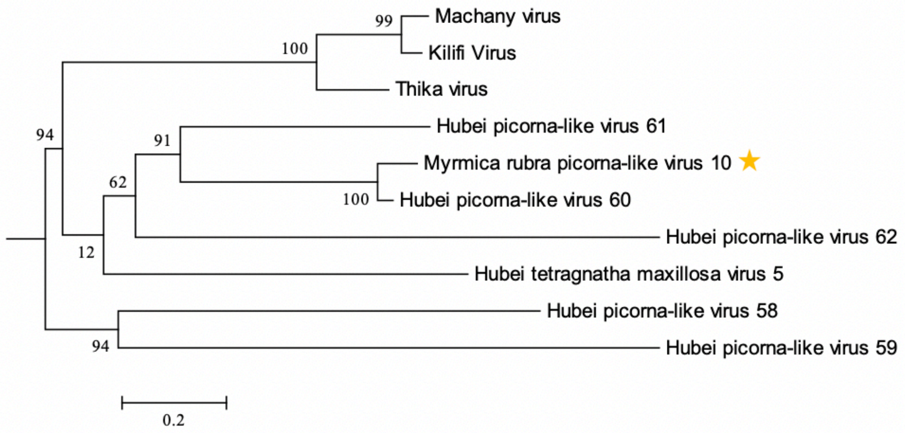
Machany virus
Kilifi Virus
Thika virus
Hubei picorna-like virus 61
Myrmica rubra picorna-like virus 10
Hubei picorna-like virus 60
Hubei picorna-like virus 62
Hubei tetragnatha maxillosa virus 5
Hubei picorna-like virus 58
Hubei picorna-like virus 59
99
100
94
91
62
100
12
94
0.2

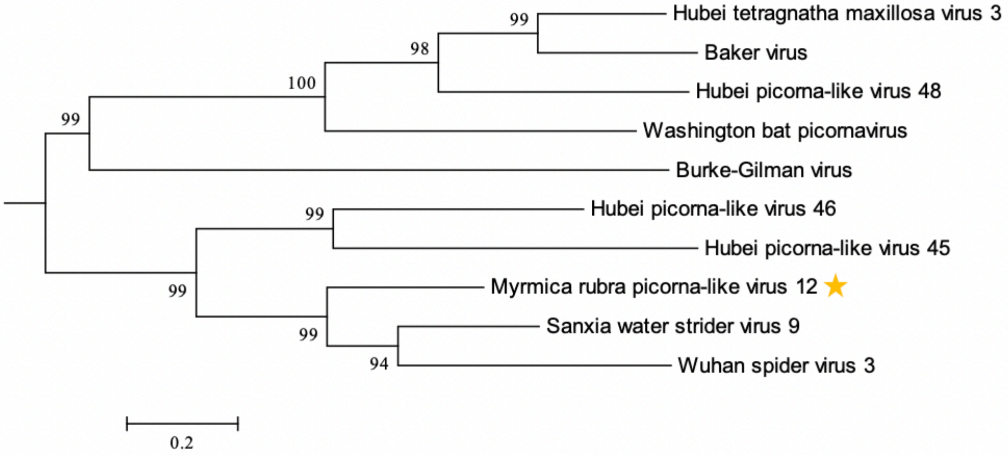
Hubei tetragnatha maxillosa virus 3
Baker virus
Hubei picorna-like virus 48
Washington bat picornavirus
Burke-Gilman virus
Hubei picorna-like virus 46
Hubei picorna-like virus 45
Myrmica rubra picorna-like virus 12
Sanxia water strider virus 9
Wuhan spider virus 3
99
98
100
99
99
99
99
94
0.2

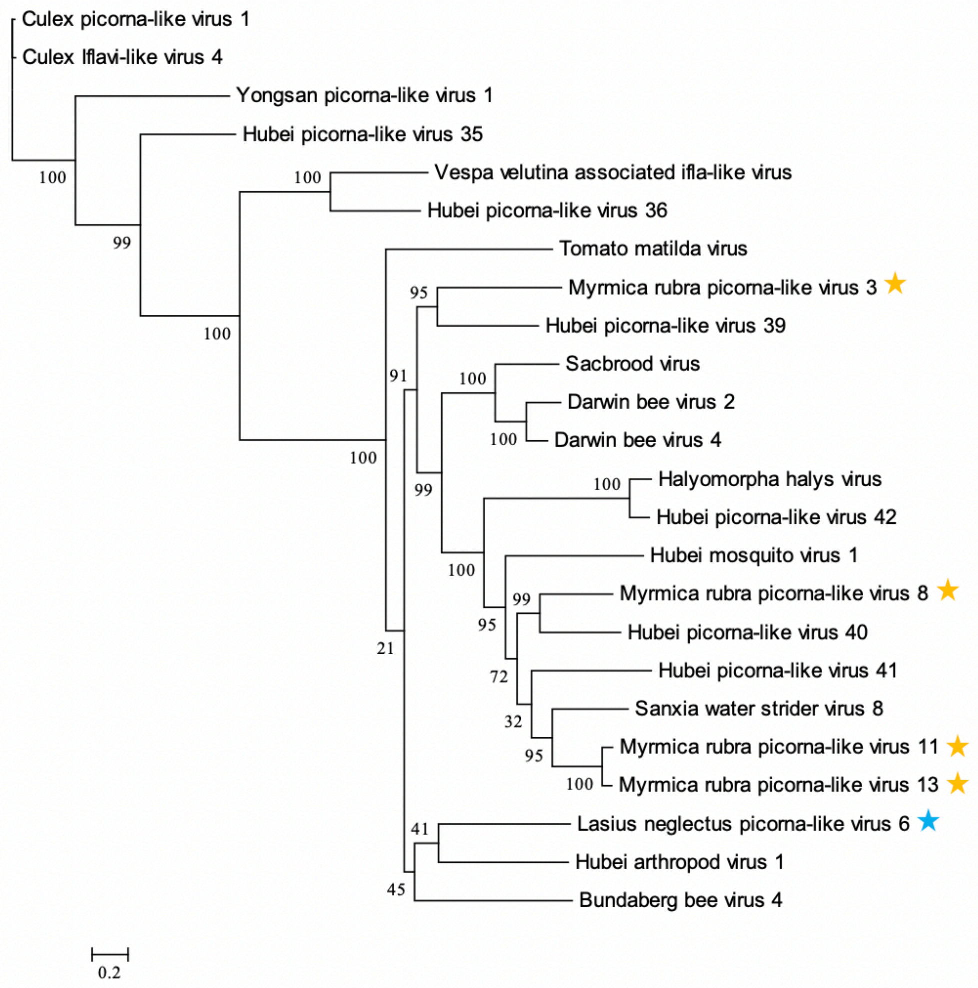


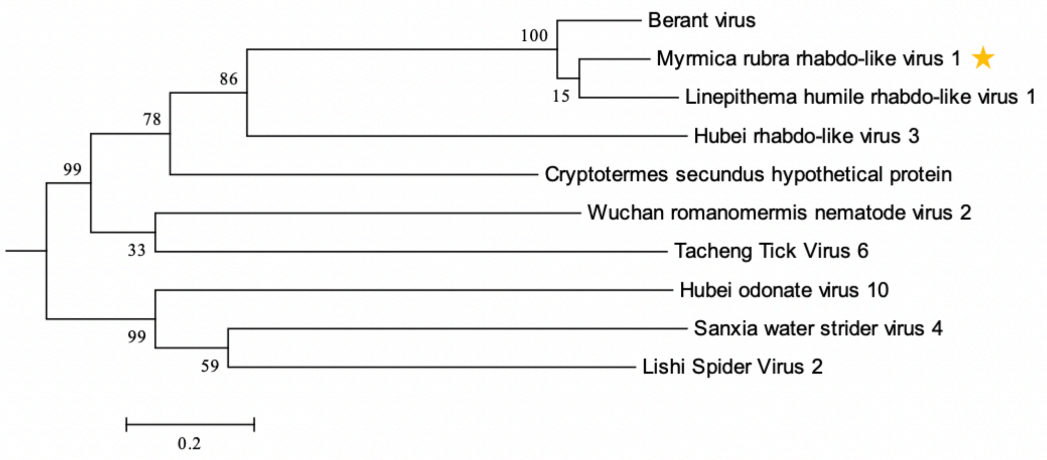


**Supplemental Figure 2. Phylogenies of the novel viruses found in this study.** Phylogenetic trees for the novel viruses discovered in this study, based on RNA dependent RNA Polymerase amino acid sequences (RdRP) of each virus and its nine best BLASTP hits. If the virus sequence matched several database viruses common to another virus of this study, their phylogenies were estimated jointly. Viruses discovered in *Li. humile* are marked with green stars, in *La. neglectus* with blue stars and in *M. rubra* with yellow stars. The branch supports of the trees are posterior probabilities based on an approximate Bayes method in PhyML v. 3.0.

**Supplemental Figure 3**

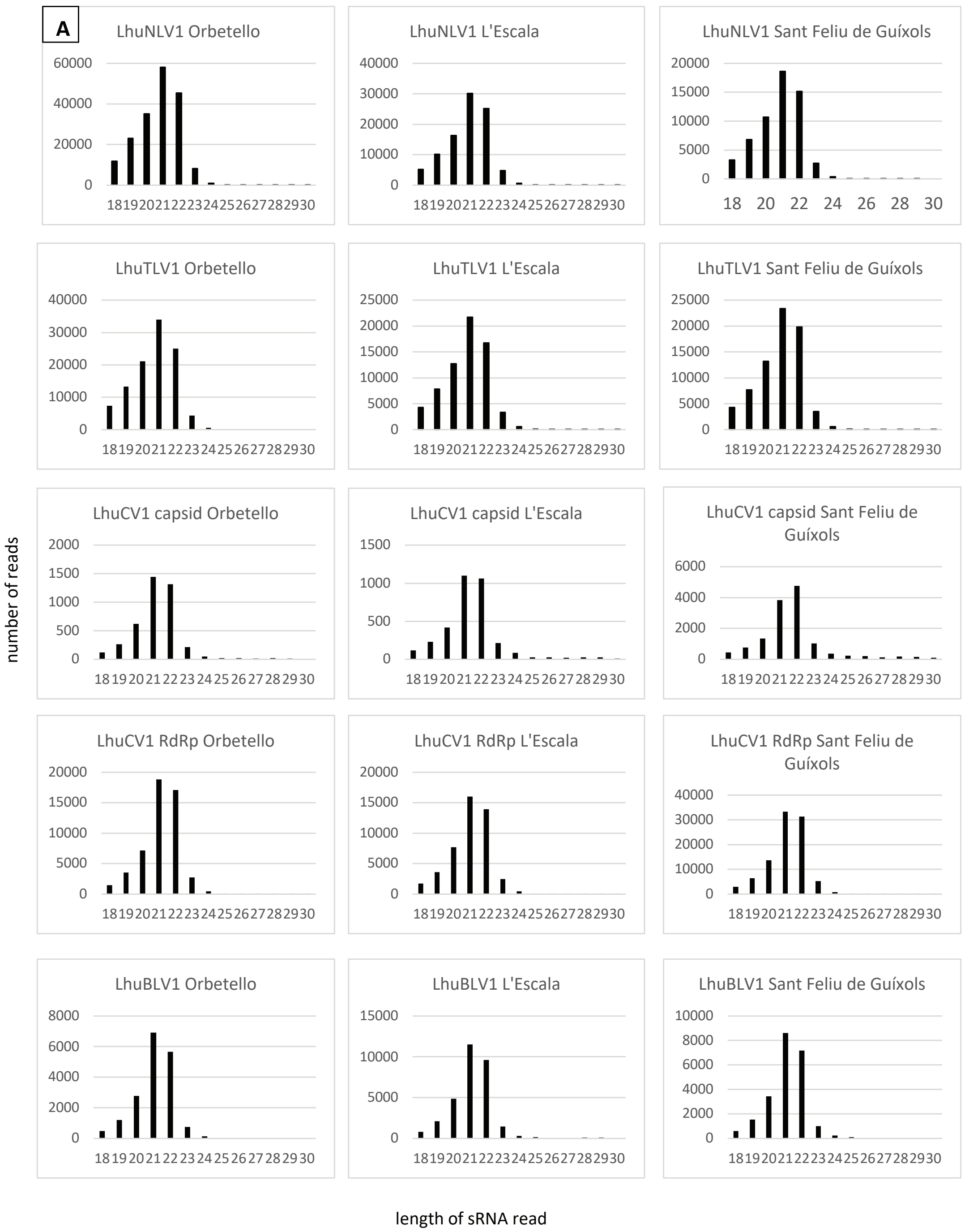

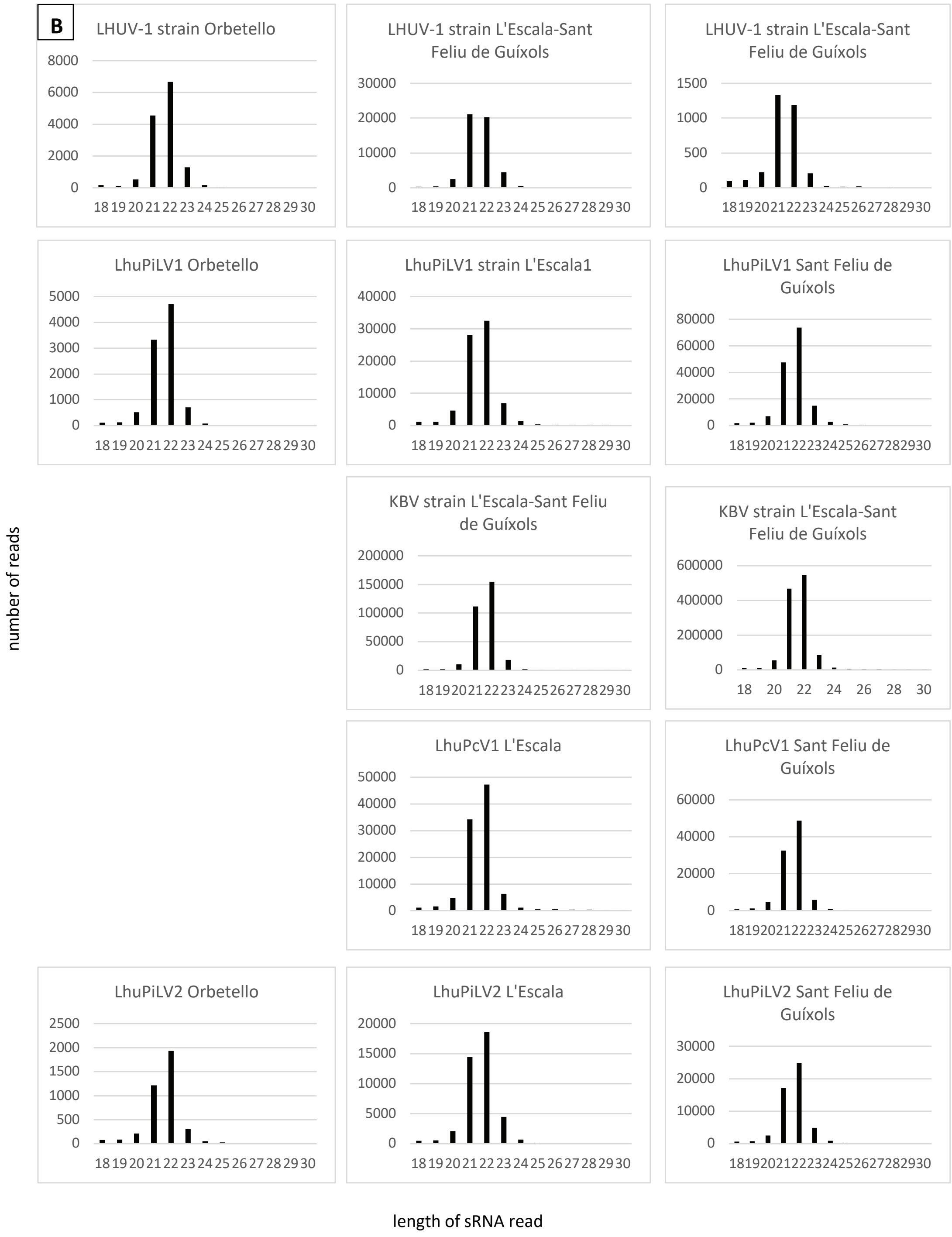


**Supplemental Figure 3. Diversity of small RNA (sRNA) size distributions in *Li. humile*.** Comparison of the sRNA size distribution of all viruses causing an RNAi response in at least two populations of *Li. humile* shows a peak mainly at 21 nt (A) or 22 nt (B) consistently across populations (populations 1: Orbetello, left; 2: L'Escala, middle; 3: Sant Feliu de Guíxols, right).

**Supplemental Figure 4**

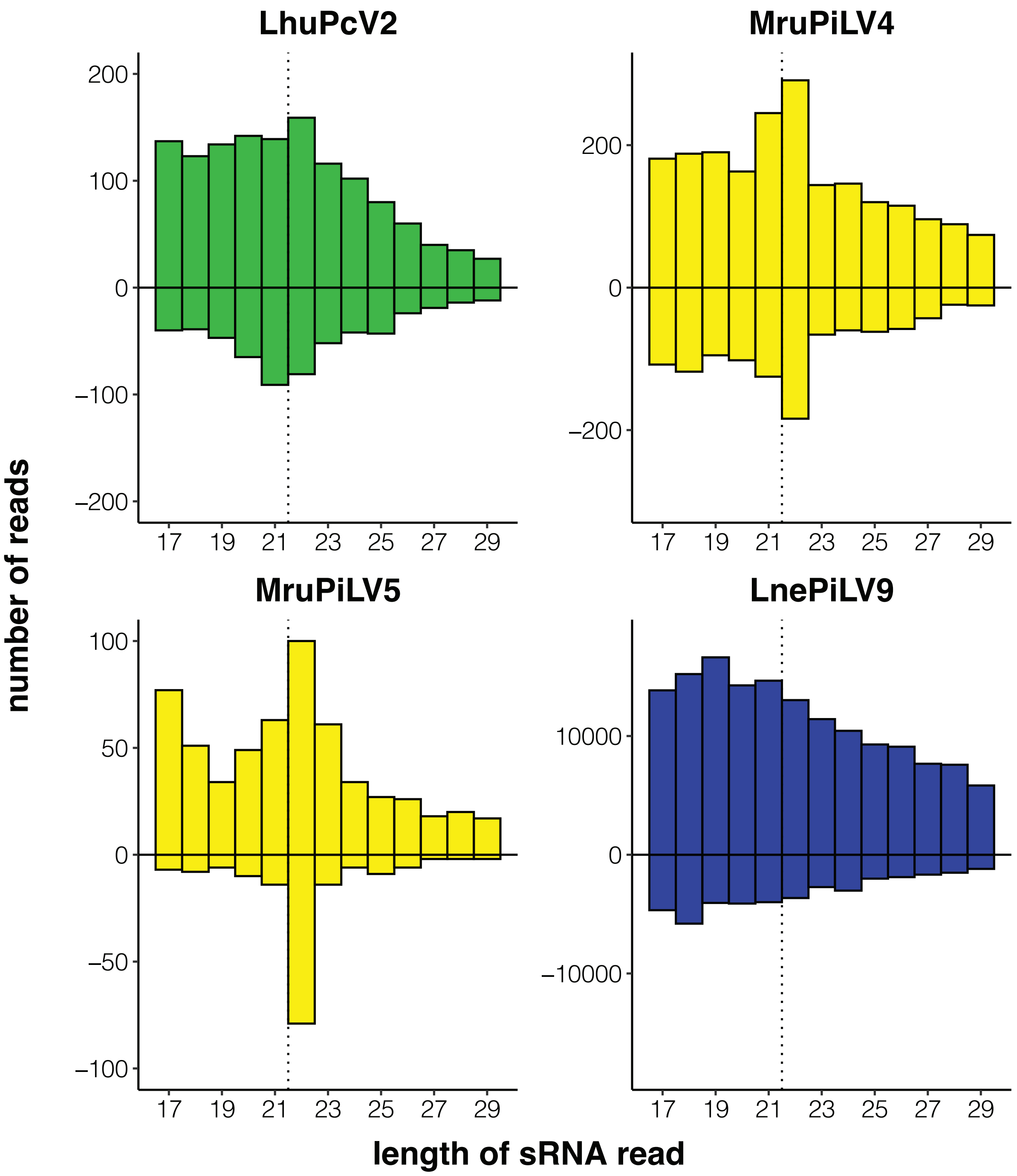


**Supplemental Figure 4. Atypical sRNA distributions.** (A) One virus infecting *Li. humile* (green) and two infecting *M. rubra* (yellow), showed atypical, broader sRNA distribution around main peaks at 21-22 nt with an overall low sRNA read count. (B) One virus infecting *La. neglectus* (blue) showed a broad sRNA size distribution in the absence of a main size class, at a high sRNA count; dotted vertical line separates 21 nt and 22 nt position. In all four cases, positive strands were overrepresented compared to negative strands.
